# Cooperative motility emerges in crowds of T cells and prevents jamming

**DOI:** 10.1101/2024.10.21.618803

**Authors:** Inge MN Wortel, Jérémy Postat, Mihaela Mihaylova, Mauricio Merino, Aanya Bhagrath, Aysha Cerf, Maryl Harris, Lin Wouters, Lucas E Wiebke, Daniel R Parisi, Judith N Mandl, Johannes Textor

**Affiliations:** Institute for Computing and Information Sciences, Radboud University, Nijmegen, the Netherlands; Medical BioSciences, Radboudumc, Nijmegen, the Netherlands; McGill Research Centre for Complex Traits, McGill University, Montréal, QC, Canada;; Department of Physiology, McGill University, Montréal, QC, Canada; Department of Microbiology and Immunology, McGill University, Montréal, QC, Canada; Instituto Tecnológico de Buenos Aires (ITBA), C.A. de Buenos Aires, Argentina; Instituto Tecnológico de Buenos Aires (ITBA), CONICET, C.A. de Buenos Aires, Argentina

## Abstract

Interacting, self-propelled particles are prone to jamming when crowded. This well-described phenomenon is shared by diverse systems including cars, animal colonies, and pedestrians. T cells, essential effectors of adaptive immunity, seemingly defy this principle: the rapid migration enabling their protective function persists even in tightly packed tissue environments – from the thymus where T cells develop, to lymphoid organs they survey for antigen, to tissues they clear from infection. Here we studied T cell crowds by combining experiments of T cells migrating in microfluidic devices with *in silico* models. We observed that while single T cells are highly heterogeneous in their motility, in crowds they synchronized their speeds and formed stable, motile trains. Our models showed that the emergence of this flocking-like behavior can be explained by a combination of two interaction mechanisms at the cell-cell interface: adhesion maintains cohesive T cell groups, and force transmission accelerates slower cells. Not all immune cells flock when they are crowded: neutrophils in the same settings slowed down with increasing cell density. Thus, cooperative motion may enable T cells to remain motile in densely packed tissue environments, preventing jams that curtail the motion of other crowded systems.

## Introduction

T cells are among the fastest immune cells in the body and their continuous motion is key to their function throughout their lifespan – from developing in the thymus to infiltrating tumors or infected tissues ^1,2^. Cell motion is also critical for the initiation of adaptive immune responses in secondary lymphoid organs (the spleen and lymph nodes), where T cell motility facilitates the highly specific interactions with dendritic cells bearing antigens ^3,4^, ensures a rapid response time to infections ^5–7^, and enables T cells to obtain the necessary signals for their survival and homeostasis ^8,9^. Intravital imaging experiments recording the movement of T cells in tissues of living mice have shown that T cells move in a random walk-like fashion guided by chemokine gradients and extracellular matrix structures ^10–15^. However, such imaging experiments usually only visualize a small fraction (< 1%) of T cells, generating the impression of T cells as autonomous agents roaming freely in empty space. Fully labelled tissue images highlight that T cells are often tightly packed with no apparent room to maneuver ^15–17^. How T cells maintain their fluid motion despite the seemingly adverse conditions of high cell density, and do so across a variety of crowded tissues, remains unclear.

Crowds of self-propelled particles – known as active matter – have been studied in different contexts: everyday scenarios such as dense vehicular traffic during rush hour, pedestrians navigating crowded train platforms on their way to work, or even grains of salt stuck in a shaker illustrate that continuous motion in a tightly packed space is not an easy task. Indeed, not only are dense crowds prone to jamming, but they can often behave in complex and unexpected ways ^18,19^. Emergent properties of active matter systems cannot usually be explained by extrapolating from the behavior of the individual agents. For instance, non-trivial crowding phenomena in the study of pedestrian dynamics include the “faster-is-slower” effect where the rushing of pedestrians to narrow exits leads to slower escape of the crowd as a whole ^20–22^. By contrast, the non-selfish behavior of ants keeps exits clear and results in more efficient egress ^23^. Despite the intuitive appeal of analogies with crowded particles, these insights may not translate to cell crowds, which differ from other active matter systems in that they are deformable and have no momentum. Pioneering work has shown that active matter principles can nevertheless apply to cells: epithelial and endothelial cell sheets jam at high cell densities, preventing cells from moving out of their home tissues; when cells manage to escape, this can have pathological consequences ^24,25^. Within such 2D cell sheets, phase transitions between fluid-state and solid-state cell collectives have been described that are density-dependent ^25–28^. Yet it remains unknown to what extent similar principles apply to T cells, which exhibit autonomous rather than collective motility ^29^, move much faster than epithelial cells, and navigate complex 3D tissues.

Combining migration experiments inspired by pedestrian dynamics with Cellular Potts simulations of T cell motility, we investigated the features of T cell crowds that result in resilience to jamming. Our work suggested that T cells self-organize into flock-like motile groups to maintain their rapid motility. In simulations, this cooperative cell motility could be explained by a combination of adhesion between cells and force transfer at cell-cell interfaces. We showed that in crowds of another rapidly moving immune cell type, the neutrophil, such cooperative motility did not emerge. Neutrophil speeds were substantially lower at increased density, suggesting that fluid motion in crowded conditions is not a universal characteristic of all crowded leukocyte populations. Overall, our work provides new insights into dynamic processes underlying immune responses and also posits T cells as interesting model systems in crowd dynamics.

## Results

### T cells maintain their motility in crowded single-lane traffic and form cell trains

We first set out to quantify the density of T cells across murine and human tissues in which they have been described to be highly motile ^30–32^. To do so, we analyzed confocal microscopy images of tissue slices from mice (thymus and lymph node) and humans (tonsil and primary melanoma in the skin). We found high-density regions of thymocytes or T cells across all tissues analyzed – with areas of large crowds of T cell aggregates (on the order of 1000-3000 T cells in a single z-plane) in the thymic cortex and medulla, the T cell zone in lymphoid organs, and even in melanoma tissue samples **(Fig. 1a)**. Thus, migrating T cells can form very dense clusters across diverse tissue types and spanning different phases of their lifecycle – from their early development to their activity in tissues during immune responses. In these dense regions, center-to-center distances between T cells revealed that ∼50% of T cells had nearest neighbor distances consistent with direct cell-to-cell contact (4-5 µm for mouse T cells and 6-7 µm for human T cells, **Fig. 1b**).

**Figure 1:**
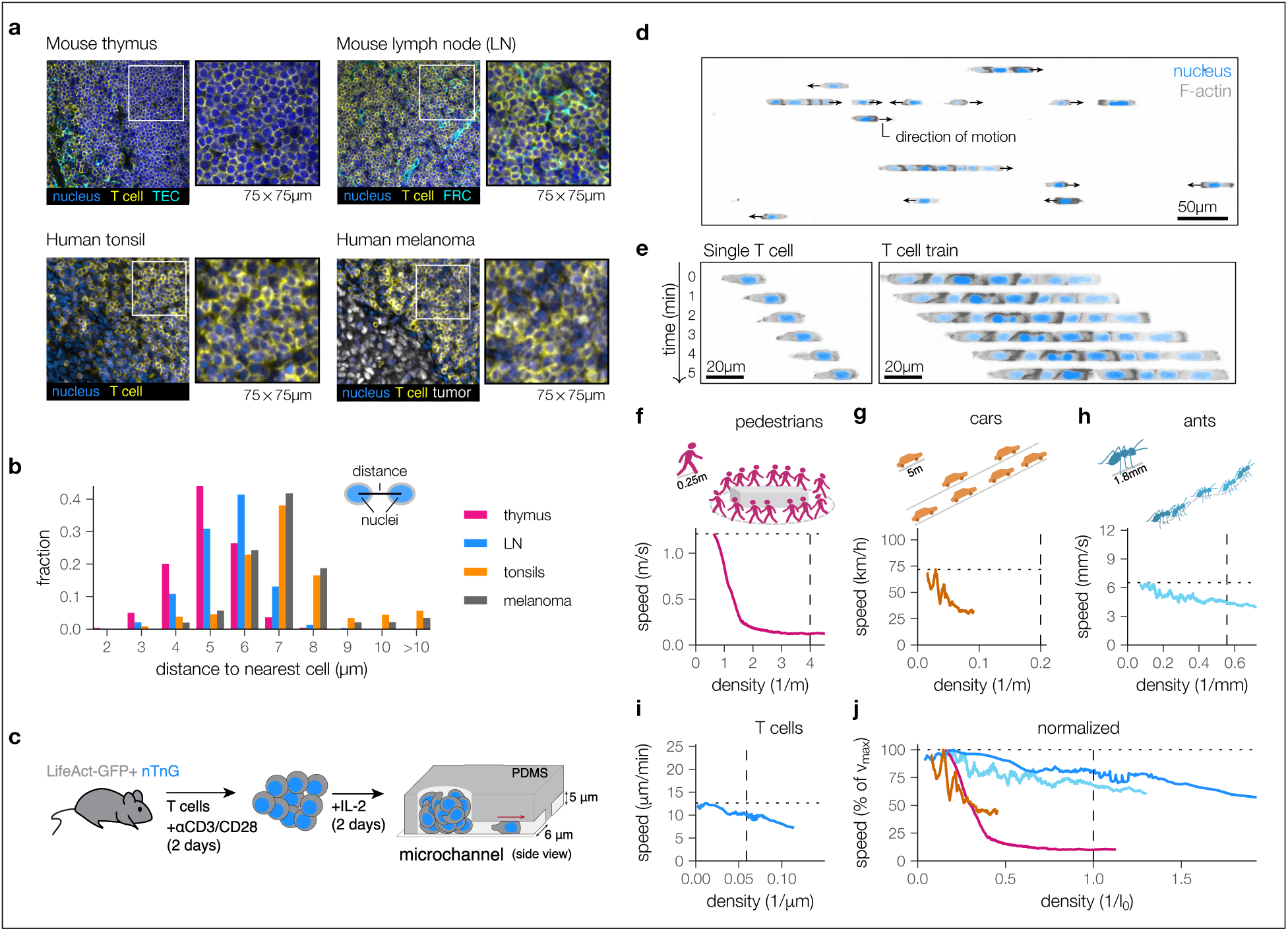
T cells remain highly motile in crowded single-lane traffic. **(a)** Representative tissue sections showing thymocytes in mouse thymus (both cortex and medulla regions), T cells in mouse inguinal lymph nodes (LN), as well as human tonsil tissue and skin melanoma. Insets show indicated areas at 2.33x magnification. Labeling for thymic epithelial cells (TEC), with keratin-5; fibroblastic reticular cells (FRC) with ER-TR7; nucleus with DAPI, and T cells in thymus with CD4 and CD8, and for LN, tonsil, melanoma with CD3. **(b)** Distances (measured between centers of nuclei) between T cells and their nearest neighbor T cells in 9 regions of interest (ROIs) per tissue (3 ROIs per specimen × 3 biological replicates). **(c)** Schematic of experimental set-up: T cells were extracted from the lymph nodes of Lifeact-GFP x nTnG mice, activated with αCD3/CD28 in the presence of IL-2, and loaded into fibronectin-coated, PDMS microchannels of 6 μm width and 5 μm height 4 days after activation. **(d,e)** Example images from the live cell imaging experiment (**Video S1**) described in (c) of T cells migrating through straight microchannels (d) and of a single T cell and a train of 7 T cells moving in a microchannel over time (e); F-actin in grey, and nucleus in blue. **(f-i)** Fundamental diagrams displaying the speed-density relation for single-lane traffic of pedestrians (f), cars (g), ants (h), and T cells moving in microchannels (i). Lines reflect the moving average of speeds measured at increasing densities, with local density measured as the inverse of the nearest-neighbor distance (raw data: Fig. **S2**). Horizontal dotted lines: maximum of the moving average speed. Vertical dashed lines: density corresponding to a nearest neighbor at distance l_0_, representing the typical agent size. (j) Fundamental diagrams from (f-i) normalized to the max speed (dotted horizontal lines) and density 1/l_0_ (dashed vertical lines). Data shown in i, j are from independent primary T cell cultures from 9 mice.

We next asked how such high cell densities impact T cell migration. While intravital microscopy has been powerful for investigating the motility patterns of T cells in various organs ^33^, in such datasets it is challenging to untangle the cell-intrinsic versus environmental constraints driving the observed migration behavior. Hence, to investigate the impact of cell density on T cell motility, we designed an experimental system that would enable us to separate the effects of T cell intrinsic properties, cell-cell interactions, and environmental constraints. We focused on single-lane traffic, the simplest system studied in crowd dynamics. This well-controlled setting has become a cornerstone in understanding emergent behaviors of crowds ^34–38^. By limiting the motion of individuals to a single lane and preventing them from overtaking each other, individuals are constrained by the speeds of their neighbors. This set-up magnifies the impact of agent-agent interactions while homogenizing environmental structure, enabling comparisons across vastly different moving particle systems.

To implement a single-lane traffic system for T cells, primary T cells were isolated from mice expressing a fluorescent reporter for filamentous actin (Lifeact-GFP) and a fluorescent tdTomato protein localized to the nucleus (nTnG), activated *in vitro* with CD3 and CD28 antibodies, and loaded into microfluidic devices fabricated from polydimethylsiloxane (PDMS) with straight channels coated with fibronectin (**Fig. 1c**). T cells in the devices moved spontaneously, without the addition of chemokine, at around 12 μm/min, and were confined to single lanes (**Fig. 1d** and **Video S1**, left). Unexpectedly, when present in channels at high cell densities, T cells frequently aggregated into rapidly moving cell trains (**Fig. 1e** and **Video S2**). The prevalence of trains depended strongly on the overall cell density, which varied between movies (**Fig. S1a**), but we found that up to 50% of the time T cells were migrating in trains of more than 5 cells (**Fig. S1b**). Because cell motility was also heterogeneous across biological replicates, even when looking only at single, non-touching cells (**Fig. S1c**), we scaled all track speeds based on the movie median of single cells before pooling the data for further analysis, allowing us to separate density-dependent effects on motility from biological variation between cells (**Fig. S1d-g**).

Next, we used these data to analyze T cell crowd dynamics. A key method for probing the dynamics in single-lane traffic is to generate the fundamental diagram of the relation between speed and local density, which depends strongly on the type of individual-level interactions. For example, pedestrians queueing in single file slow down as individuals get closer to each other, as do cars in dense traffic (**Fig. 1f,g**, and **Fig. S2a,b**) ^34–36,39^. By contrast, ants are a crowded system of non-selfish, social insects that stays motile at high densities ^37^ (**Fig. 1h** and **Fig. S2c**). Interestingly, T cells in straight microchannels maintained a relatively constant cell speed across varying densities (**Fig. 1i**). To directly compare the fundamental diagrams of these four different motile particle systems from experimental data, we normalized agent speeds and densities such that a local density of 1 was achieved at the average distance where individual agents touch. Pedestrians and cars reduced their speed at densities substantially below the direct-contact threshold (**Fig. 1j**). For T cells, there was no clear transition point at which they slow down – owing in part to their highly variable sizes (**Fig. S2d**) – but the diagram was mostly flat until a normalized density of around 0.5 where the largest cells touched (**Fig. 1i,j** and **Fig. S2e,f**).

In summary, we observed that T cells exhibit single-lane crowd dynamics akin to those of ants, which are known for their resilience to jamming. T cells maintained their speed in high-density regions and self-organized into highly motile trains.

### The motility of individual T cells in single-lane traffic is highly heterogeneous

To understand the interaction mechanisms underlying the flocking-like motility of T cell trains, we first set out to characterize the baseline behavior of cells that were not in trains. We isolated single-cell (sc) tracks of cells traveling without touching other cells for a duration of at least five minutes (**Fig. 2a**, **Video S1**). Across 1481 normalized tracks from 9 movies, we extracted 544 sc-tracks, and found that individual T cells moving in straight channels exhibited a striking heterogeneity in motility. One source of variation was cell speed, with average sc-track speeds of different cells ranging from less than 5 μm/min to more than 20 μm/min (**Fig. 2b**). In addition, the speed of individual cells varied over time, as shown by the standard deviation in instantaneous speeds observed for individual sc-tracks (**Fig. 2c**). Cells varied considerably in the time that they spent in a non-motile state, with some cells frequently alternating between “stop”, “go”, or “turn” states without contact with other cells (**Fig. 2d**, left), while others moved steadily in one direction despite speed fluctuations (**Fig. 2d**, right). To quantify T cell stopping and turning behavior, we classified cells as “stopped” when the magnitude of their instantaneous velocity vector was below 5 μm/min, and as “turned” whenever this vector reversed direction. Accounting for the inherent noise in the cell tracks using Kalman smoothing, we extracted cell stops and turns (**Fig. 2e**, **Video S3**), and found that the arrest coefficient (percentage of time spent in the stop-state) was highly heterogeneous: while 47% of cells did not stop for the entire duration they were tracked, others had arrest coefficients between 0 and 50% (**Fig. 2f**). This effect was separate from the heterogeneity in speed, as sc-track speeds still varied between 5 μm/min and 30 μm/min even when “stop” intervals were not considered.

**Figure 2:**
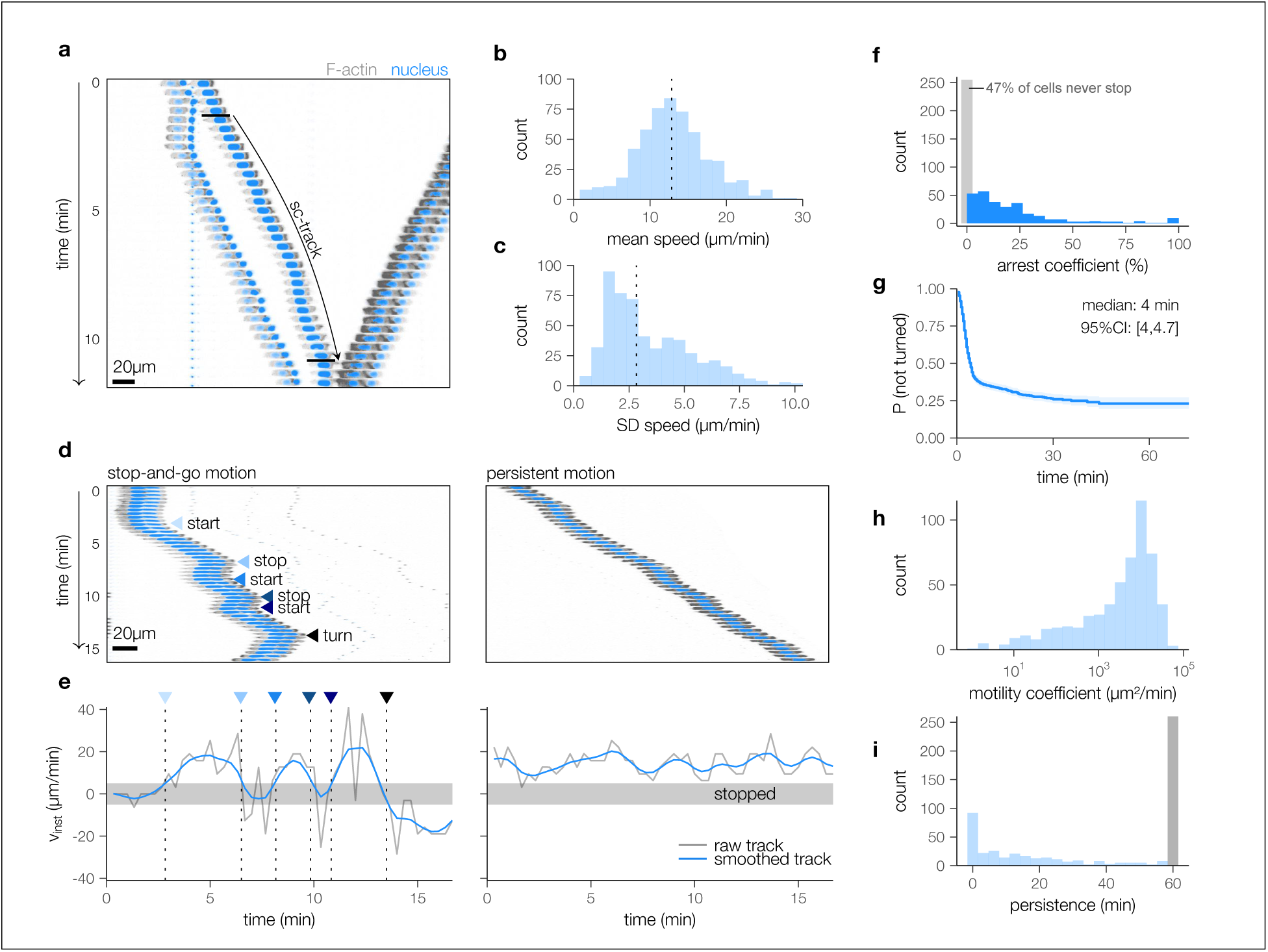
The motility of single T cells in microchannels is highly heterogeneous. **(a)** Single cell track example (sc-track, indicated with arrow) extracted from the microchannel live cell imaging data. **(b,c)** Distribution of the mean (b) and standard deviation, SD (c), of instantaneous T cell speeds for each sc-track. **(d)** T cell kymograph examples showing a cell with a stop-start behavior (left), and a continuously moving cell with fluctuating speed (right). **(e)** Raw (gray) and smoothed (blue) directed instantaneous velocity (v_inst_) for the kymographs shown in (d). Cells are considered stopped when smoothed |v_inst_| < 5 µm/min (gray region). **(f)** Distribution of arrest coefficients (percentage of time spent stopped) in sc-tracks. **(g)** Kaplan-Meier plot of persistence of intervals in which a cell does not turn or stop. **(h,i)** Motility coefficients (h) and persistence times (i) of sc-tracks; gray bar, persistence time of 60 minutes or more. Data in b, d-g are based on 544 normalized sc-tracks pooled from 9 movies, taken in 3 independent experiments, where each movie uses a T cell culture taken from an independent mouse.

Lastly, cells varied in the duration of persistent motion without stopping or turning. We assessed the duration of the persistent state with a Kaplan-Meier analysis to account for cells leaving the field of view (**Fig. 2g**). About 1 in 4 sc-tracks were persistent over half an hour to an hour without any observed stopping or turning. Altogether, the variability in speed, stopping, and turning behavior translated into variable motility coefficients and persistence times between individual T cells (**Fig. 2h,i**). Thus, our data showed that cell-intrinsic heterogeneity was a defining feature of T cell motion in simple, obstacle-free microchannels — raising the question of how such heterogeneously moving cells would be able to form motile trains.

### Colliding T cells form cohesive trains and match their speeds with neighboring cells

The T cell trains we observed (**Fig. 1d,e** and **Video S2**) were reminiscent of animal collectives where individuals synchronize their motion, such as flocks of starlings or schools of fish ^40,41^. Such higher-order global behaviors of animal collectives can be explained by simple interaction rules between individuals ^42,43^. Following Reynolds ^42,44^, two critical factors that underlie flock formation are: *cohesion*, a force that prevents nearby agents from drifting apart (but leaves enough room for maneuvering); and *velocity matching* whereby neighboring agents synchronize their speeds and directions.

To investigate whether the flocking-like behavior exhibited by T cells in microchannels can be explained by similar principles, we tested for evidence of cohesion and velocity matching. We first focused on the simplest cell-cell interactions in single-lane motility: collisions between two cells (**Fig. 3a**). We tracked the outcomes of a total of 698 collisions in the dataset and found that in most cases, colliding cells remained close (**Fig. 3b**) and within 5 minutes post-collision, the cells quickly recaptured most of their initial speed (**Fig. 3c**). A Kaplan-Meier analysis accounting for censoring due to cells leaving the observation window, or cell pairs being joined by other cells, indicated that almost 75% of one-on-one colliding T cells remained together as pairs for the time window they were within the field of view (**Fig. 3d**). While longer T cell trains did lose some adjoining cells more quickly, almost half of the trains of >4 cells remained together for 30 minutes or longer (**Fig. 3d**). Many of the cohesive cell trains were highly motile but also had large standard deviations in speed over time due to frequent pauses in motility, changes in direction, or both (**Fig. 3e**). Importantly, when we examined the cell trains with the greatest standard deviation in speed in more detail, we found that even during turns or changes in speed, cells in a train preserved their matched directions (**Fig. 3f**). These data suggested that T cells have a strong tendency to maintain contact for extended periods of time.

**Figure 3:**
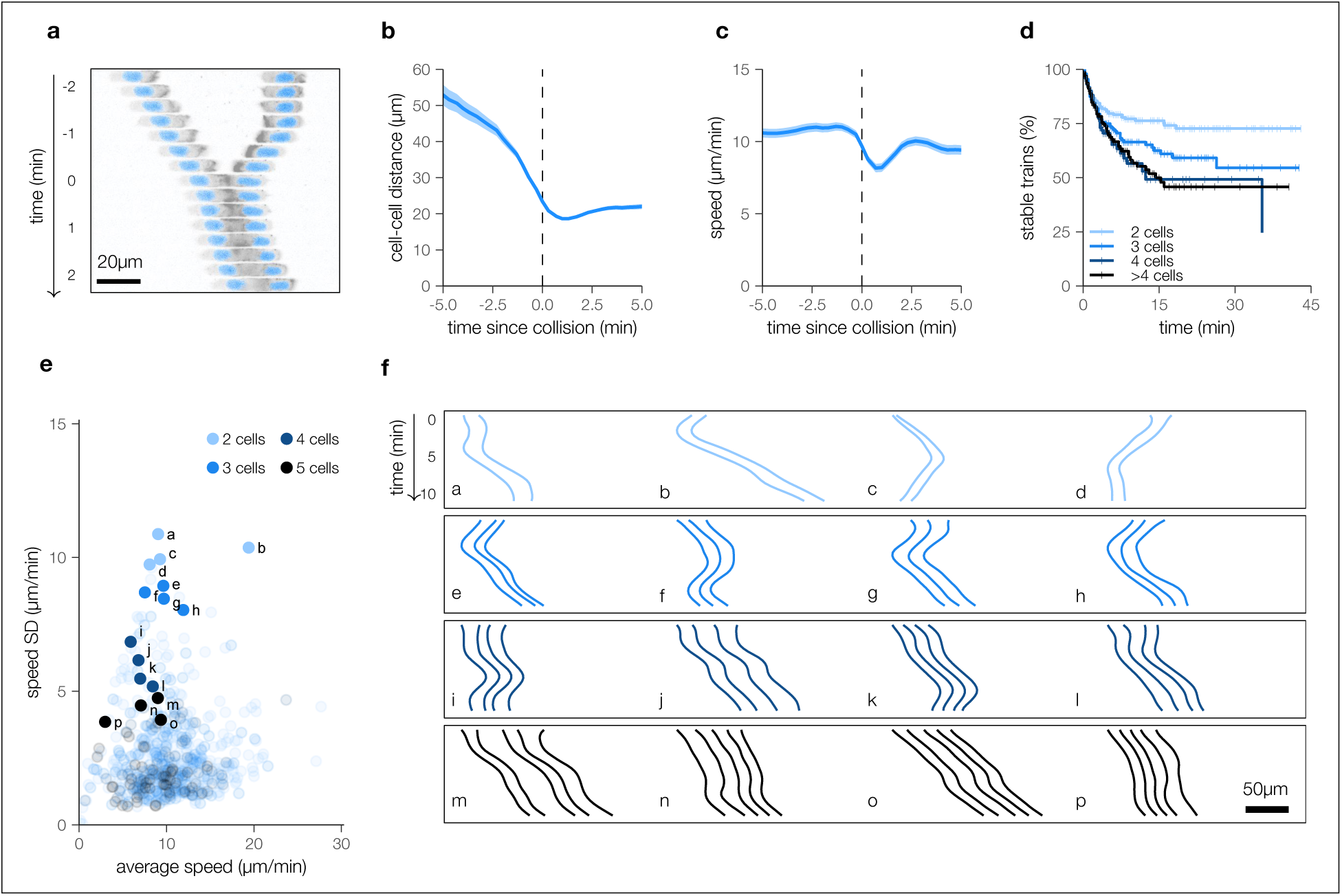
T cell trains in single-lane traffic are motile and cohesive. **(a)** Example of two single cells forming a train of size 2. **(b,c)** Distance (b) and speed (c) between cells in a pair before and after collision; solid lines: mean, shading: standard error based on N=316 collision events. **(d)** Kaplan-Meier analysis of how long trains remain together after forming, stratified by train size. **(e)** Average speed and its standard deviation for each motile train. **(f)** Example T cell trains showing that trains can collectively take turns or make brief stops. Each T cell train shown corresponds to one of the numbered points in (e).

To test whether, in addition to their directional alignment, T cells also match their speeds to each other as observed in animal flocks ^40^, we studied the outcomes of “head-to-tail” collisions in our data, in which a faster “rear” cell bumps into a slower “front” cell (**Fig. 4a** and **Video S4**). We identified 77 head-to-tail collision events where cells remained together for at least 3 minutes afterwards. As observed for individual cell tracks, we found substantial heterogeneity in cell speed before as well as after collision (**Fig. 4b**). Strikingly, however, the front and rear cells matched their speeds post-collision (**Fig. 4c**). The matched speeds did not simply result from the rear cell “queuing up” behind the front cell, but instead, were the result of both a decreasing speed of the rear cell *and* the increasing speed of the front cell. Thus, while most (71%) rear cells slowed down after collision, front cells sped up by an average 32% over that same interval (**Fig. 4c**). In line with the collision-induced acceleration of slower front cells, the speed of T cell trains made up of >4 cells was only ∼20% lower on average than those of individual cells (**Fig. 4d**). This finding was not consistent with expectations derived from a “queuing model” where the speed of each train is dictated by the cell at the front (which must be the slowest cell in the train, which would otherwise break up; **Fig. 4e**). Such a queuing model also could not explain the many instances of trains moving rapidly in synchrony (**Fig. 4f**). We further constructed an alternative model we termed the “extended queuing model” (Supplementary Methods, and **Video S5**) that, unlike the simple queuing model, reproduces both the speed and turning behavior of sc-tracks (**Fig. S3a,b**) but in which cells do not match their speeds or have a preference to remain together. In simulations matched to the densities observed in our imaging data (**Fig. S3c**), this model was unable to explain the stability of T cell trains (**Fig. 3d**) as simulated 2-cell trains and longer trains fell apart much more rapidly than observed in our data (**Fig. S3d**).

**Figure 4:**
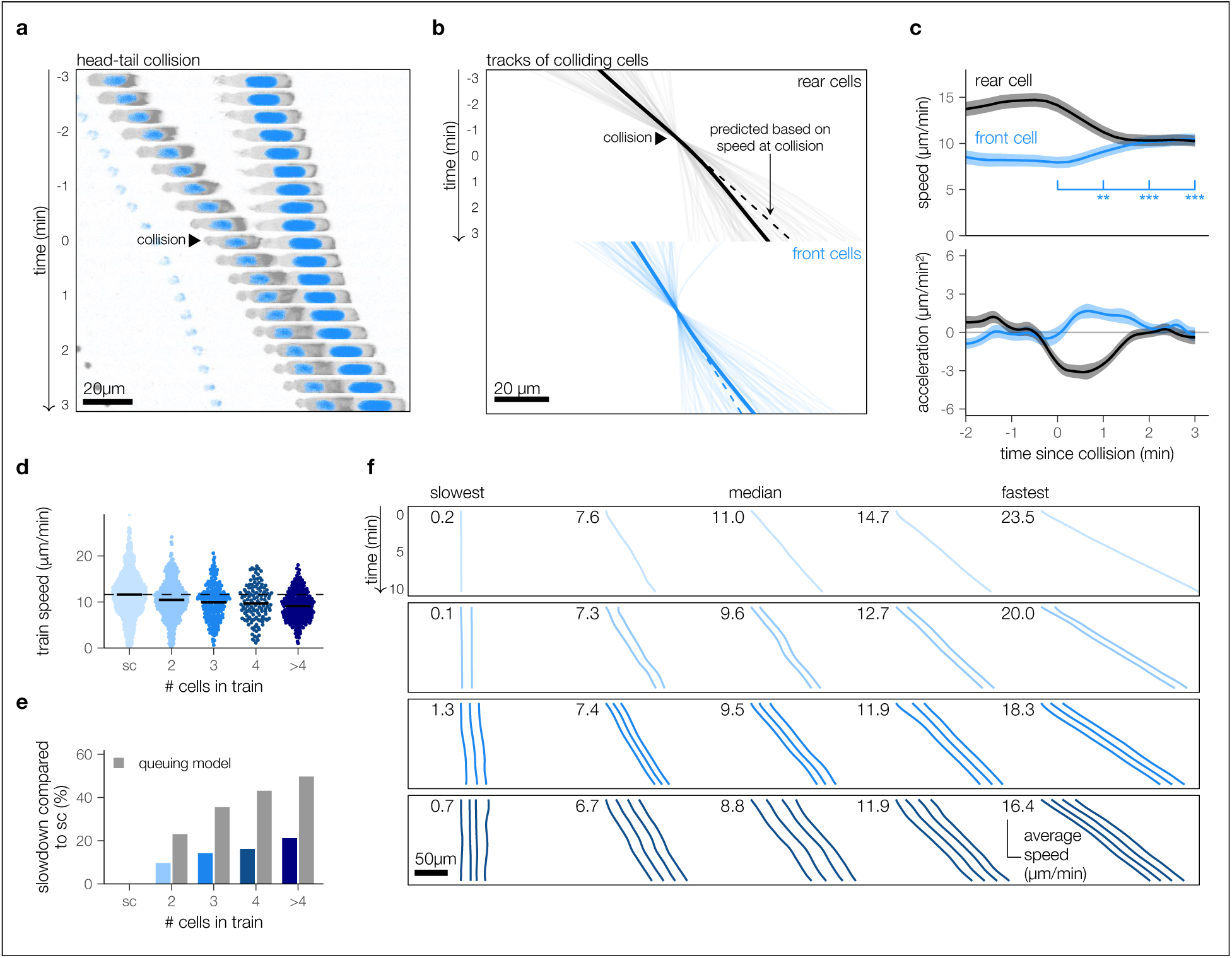
Neighboring T cells align their speeds after head-to-tail collisions. **(a)** Example of a head-to-tail collision where a faster-moving rear cell collides with a slower-moving front cell, leading to speedup of the front cell and slowdown of the rear cell. **(b)** Tracks of the rear (black) and front (blue) cells in all head-to-tail collisions observed in our data (N=77). Solid lines are average position of cells over time. Dashed lines show predicted average position based on speed at time of collision. **(c)** Speed and acceleration of individual cells estimated from smoothed cell tracks before and after time of collision (set at time = 0). Lines indicate means; shading, standard error. **(d)** Distribution of average cell speeds moving alone or in trains of varying lengths. Lines, medians; dotted line, median speed of sc-tracks. **(e)** Speed slowdown of trains (percent decrease in median speed compared to single cells) calculated from sampling the sc-track speed distribution, assuming that the train speed corresponds to the lowest sampled speed (expected train speed from queuing model, grey bars) compared to the measured actual slowdowns (blue bars). **(f)** Representative example tracks representing the speed distribution (minimum, maximum, and first to third quartiles of average speed over a 10-minute interval) of T cells moving alone, or in trains of length 2, 3 and 4 (by row).

### Local interactions at cell-cell interfaces suffice to explain T cell flocking

In animal collectives, synchronization of speed and direction is achievable by local communication between individuals. Cells can communicate via diffusing chemical signals or by mechanisms involving direct physical contact between cells. We did not find any evidence for communication without direct contact: there was no dependence of speed on cell-cell distance detected in the fundamental diagram (**Fig. S4**) and there was no apparent dependence of directionality on distance either (**Fig. S4**). Therefore, we focused on investigating whether the observed T cell cohesion and alignment could instead result from local forces generated at the cell-cell interface. We performed *in silico* simulations using a Cellular Potts Model (CPM) of cell migration (**Fig. 5a**) ^45–47^, with which we had previously simulated T cell migration in microchannels ^48^. The CPM represents cells as collections of pixels on a grid that can extend or retract with probabilities P_out_ or P_in_, respectively. These probabilities depend on an associated Hamiltonian energy difference ΔH, which consists of several terms including a dedicated term ΔH_act_ that models active protrusions ^47^ (Methods). Since ΔH equals the (negative) work W and CPM changes operate over a distance Δx = 1 pixel, energy difference terms |ΔH| = |W| = |F Δx| can be interpreted as force magnitudes as well.

**Figure 5:**
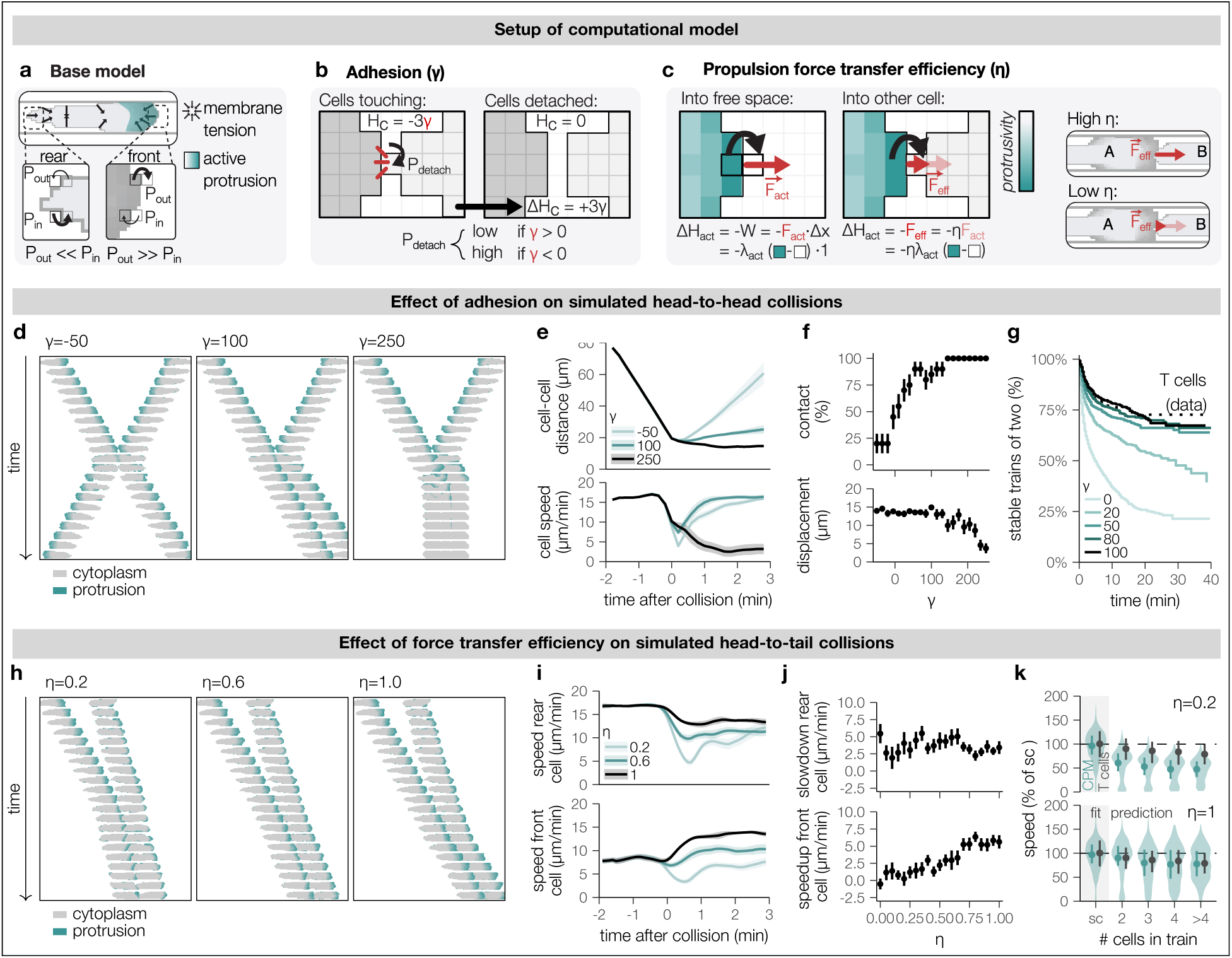
Local interactions at cell-cell interfaces generate cohesion and alignment in a computational model. **(a)** Modeling cell motility with the cellular Potts model (CPM) in which cells are represented as pixels on a grid that can be stochastically extended or retracted. In the base single-cell model, the extension and retraction probabilities (P_out_ and P_in_) depend on the local balance between membrane tension and the force generated by protrusions. **(b,c)** Specific modifications incorporated into the CPM to investigate the impact of two parameters on T cell train formation: the adhesion parameter γ representing the (change in) contact energy (Δ)H_c_ associated with separating versus maintaining a cell-cell interface (b), and the efficiency η at which protrusion-generated forces are transferred from one cell to another (c). **(d)** Example simulations of head-to-head collisions of two cells in a microchannel generated by the CPM for negative, moderately positive, and high γ. **(e)** Distances between cells and cell speeds before and after collision for negative, moderately positive, and high γ. **(f)** Percentage of cells still in contact, and displacement of cells from the position of contact 5 minutes after the time of contact, across a range of γ. **(g)** Longer-term train survival predicted by a detailed simulation based on observed cell positions and speed distributions in the microchannel data for a range of positive γ values compared to the T cell data. **(h)** Example CPM simulations of head-to-tail collisions for low, medium, and high η. **(i)** Speed of the rear and front cells for low, medium, and high η. **(j)** Slowdown of the rear cell and speedup of the front cell, measured by comparing the instantaneous speed 3 minutes after and 2 minutes before collision, across the range of η. **(k)** Train speed by train size at maximal γ compared to the T cell data predicted by detailed simulation of the microchannel data for medium and high values of η. In (e), (f), (i), and (j), lines or points indicate means and shading or bars indicate standard error of the mean for 20 simulations per parameter combination.

We hypothesized that two local factors were important to explain the emergence of stable motile trains. First, we expected adhesion between cells to stabilize the contacts between T cells, as this would prevent fast cells from detaching and moving away from slower cells. Second, for cells moving in the same direction, we reasoned that the forces generated by actin polymerization of the rear cell could lower the force required for retracting the rear of the front cell – presumably leading to an alignment of speed, or more specifically, to an increase in speed of slower moving front cells. Therefore, to test whether adhesion and force transmission would suffice to explain the emergence of T cell flocking behavior, we developed an extended CPM where we could tune both an adhesion parameter γ to determine the contact energy H_c_ between cells (**Fig. 5b**), and an efficiency parameter η that controls which fraction of the protrusion force F_act_ is transferred to a neighboring cell (**Fig. 5c**).

To investigate the adhesion and protrusion force parameters, we first fitted the two main parameters of individual cell motility in our CPM (**Fig. S5a**) to reproduce the observed speed distributions (**Fig. S5b**) and turning behavior (**Fig. S5c,d**) of our sc-tracks, and generated simulations matching the cell densities observed from the imaging data (**Fig. S5e, Video S6**). We then simulated head-to-head collisions between two cells to examine the consequences of the adhesion parameter γ on cohesion (**Fig. 5d**). In these simulations, a moderately positive level of adhesion (e.g., γ=50) was required to reproduce the observed cohesion between cell pairs: negative adhesion between cells (e.g., γ=-100) led to rapid contact dissolution, whereas excessive adhesion (e.g., γ=200) slowed or even stopped the motion of the cell pairs (**Fig. 5e**). Therefore, a trade-off between train stability and motility emerged, requiring moderately positive adhesion to achieve a high chance to form T cell pairs that remain in contact but still move rapidly (**Fig. 5f**). Similarly, in more detailed simulations where we mimicked the positions and speed distributions of the T cells in our imaging data, moderately positive adhesion was required to generate trains that remained stable for more than 30 minutes (**Fig. 5g**).

Next, we used the CPM to simulate head-to-tail collisions to investigate to what extent protrusion force transmission from the rear to the front cell led to speed matching (**Fig. 5h**). Without any force transfer, both cells involved in the collision slowed down upon contact (**Fig. 5i**), and a high force transfer efficiency was required to speed up the front cell upon collision (**Fig. 5j**). As observed in the experimental data, rear cells slowed down following contact; force transmission promoted trailing-edge retraction in the front cell but did not enhance leading-edge protrusion in the rear cell. In detailed simulations mimicking the observed T cell positions and speed distributions, high force transmission efficiency was required to reproduce the speed of longer T cell trains observed in the experimental data (**Fig. 5k**). This force transmission between T cells is consistent with the fact that T cell motility does not rely on focal adhesions, which would otherwise be expected to impose substantially greater resistance.

Taken together, in the computational model, inter-cellular adhesion and efficient protrusion force transmission through the cell-cell interface were sufficient to generate cohesion and speed matching between neighboring T cells, explaining both the stability and motility of emergent cell trains.

### Cooperative motility does not emerge in neutrophil crowds

The speed T cells can achieve is second only to neutrophils, cells of the innate immune system which are usually the first to arrive at sites of tissue injury or infection ^49–52^. Although T cells and neutrophils share an amoeboid-like migration mode that sets them apart from fibroblasts, epithelial or endothelial cells, migratory behaviors still differ substantially between T cells and neutrophils *in vivo*, likely due to their distinct functions in an immune response. While T cells need to scan for antigen, either for surveillance purposes or to trigger their effector functions as they move through dense crowds, neutrophils swarm to their destination, where they often aggregate at high cell densities and reduce their motility ^52,53^. Given our findings that T cell crowds in single-lane traffic form flock-like collectives, we asked whether this phenomenon was specific to T cells or might be generalizable to other rapidly moving leukocytes.

To address this question, we isolated mouse neutrophils from bone marrow, pulsed them with lipopolysaccharide (LPS) *in vitro*, and loaded cells into microchannels, again without addition of exogenous chemokine (**Fig. 6a**). As with activated T cells, stimulated neutrophils exhibited spontaneous and heterogeneous motility in single-lane channels (**Fig. 6b** and **Video S7**). We tracked 1380 neutrophils across 7 live cell microchannel movies and extracted 1427 normalized sc-tracks using the same procedure as previously for the T cells (**Fig. S6**). The motility of single neutrophils was similar to that of single T cells in terms of speed, arrest coefficient, and persistent directionality (**Fig. 6c**). Interestingly, in the crowded regions within the microchannels, the behavior of neutrophil trains was starkly different compared to T cells: longer neutrophil trains were much slower than shorter ones (**Fig. 6d**), and we observed several instances of jammed groups of cells that were immotile (**Fig. 6e**, **Video S8**). Notably, this was not due to dying neutrophils at the center of cell trains attracting additional cells in a feed-forward loop as has been described in neutrophils swarming *in* _vivo_ ^51,53,54^.

**Figure 6:**
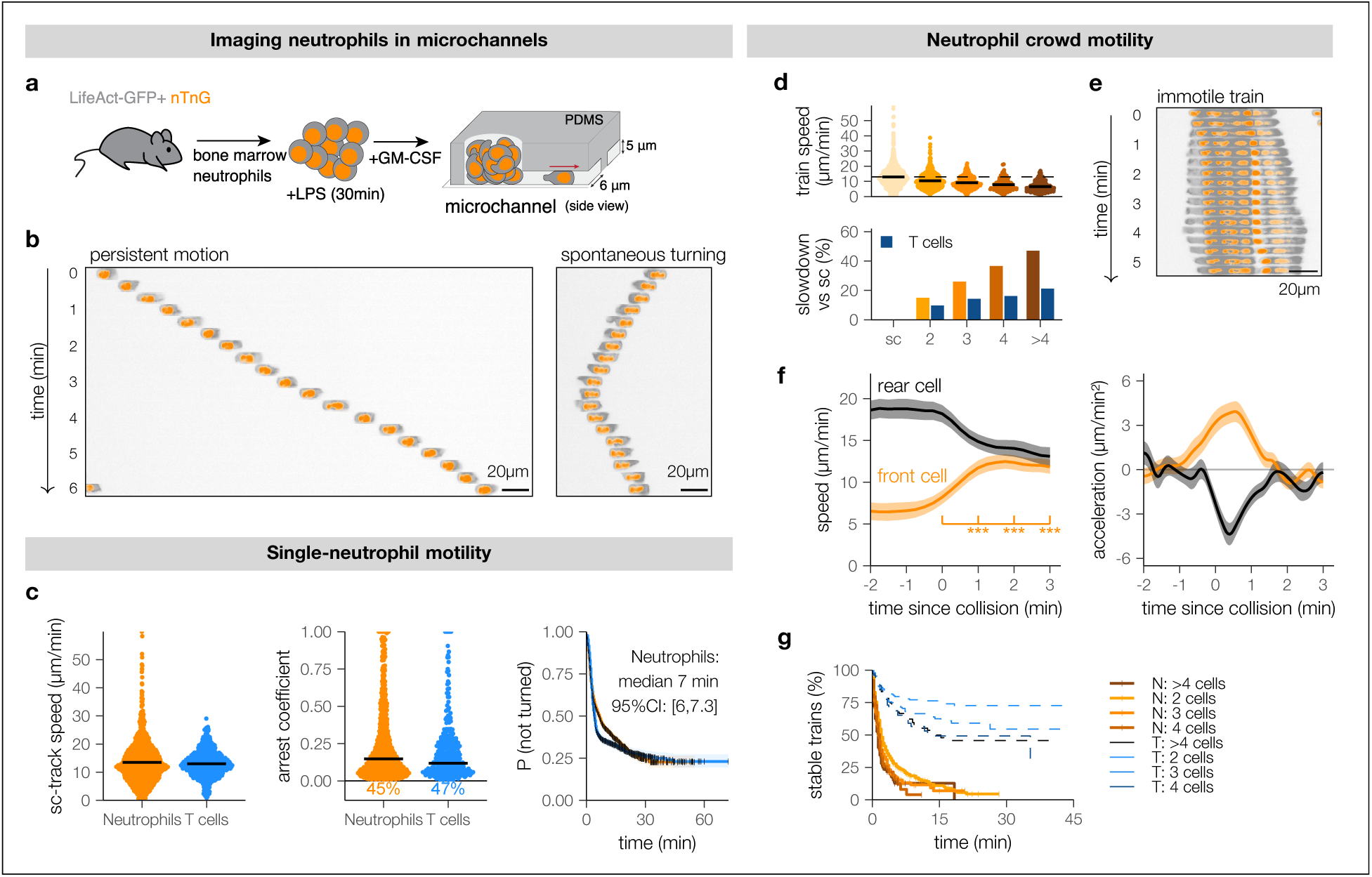
Neutrophils in single-lane traffic show alignment but not cohesion. **(a)** Schematic of experimental set-up: Neutrophils were extracted from the bone marrow of Lifeact-GFP x nTnG mice, pulsed with LPS, and loaded into microchannels as for T cells (Fig. 1). **(b)** Two examples to illustrate the heterogeneous motility behaviors of neutrophils in microchannels, with some cells moving persistently and others spontaneously turning. **(c)** Sc-tracks were extracted from neutrophil live imaging data to obtain track speeds, arrest coefficients, and Kaplan-Meier plots of persistence of intervals during which a cell does not turn, comparing them to sc-track data from T cells (from Fig. 2). **(d)** Speeds of neutrophil trains of different sizes (top) and comparison of the slowdown with train size to the T cell data (bottom; T cell data from Fig. 4). **(e)** Representative example of a neutrophil “traffic jam” (non-motile train of five touching cells). **(f)** Speed and acceleration of neutrophils involved in head-to-tail collisions (mean and standard error, N=74 collisions). **(g)** Cohesion of neutrophil (N) trains of different sizes compared to T cells trains (T) (dashed, data repeated from Fig. 3d). Panels c,e,f are based on pooled, normalized sc-tracks from 7 replicate movies.

A potential explanation for the jammed neutrophil trains was the absence of speed matching, which would make slow neutrophils unable to speed up when pairing with faster cells catching up from the rear. Indeed, the slowdown of longer trains predicted from the simple queuing model without speed matching fitted the neutrophil data well (**Fig. S7**). However, from analyses of head-to-tail collisions we found that front neutrophils in fact almost doubled their speed upon collision (**Fig. 6f**). Instead, neutrophils differed from T cells in their cohesion: unlike T cells, most neutrophils did not remain together for extended periods of time (**Fig. 6g**). Faster neutrophils running into slower cells in many cases turned away, leaving slow cells to form groups with each other – ultimately creating a selection bias whereby longer trains tended to be constituted of slower cells. Interestingly, these differences in crowd dynamics between T cells and neutrophils were not nearly as obvious in the fundamental diagram (**Fig. S8**), which considers only the distance to the nearest neighbor rather than the number of close-by neighbors. This suggests that the fundamental diagram may fail to capture relevant crowd dynamics that follow from direct contact rather than interactions over longer ranges.

Thus, while activated T cells displayed a resilience to cell jamming, this was not an intrinsic feature of all leukocytes: We found that the flocking behavior of T cells was not recapitulated in migrating neutrophils which, while able to match their speeds, did not demonstrate cohesion.

## Discussion

The first intravital microscopy studies of T cells highlighted their rapid motility in physiological contexts ^11–13^ and subsequent work has emphasized the critical reliance of immune function on T cell motion ^6,7^. Distinct T cell motility behaviors have been described *in vivo*: from the random walk-like, stop-and-go motion of naive T cells in lymph nodes ^13,55,56^, to directed motion in positively selected thymocytes ^57^, and superdiffusive motion of effector T cells in the brain ^58^. Even within the same context, motility can vary widely between cells and over time ^13,59–61^. Yet it has been difficult to assess how much of this variation is a function of environmental structure versus cell-intrinsic properties. Whether differences in motility reflect interactions with physical obstacles ^10,62^, chemokines ^55^ or peptide-MHC ^30,63,64^, or arise from cell-cell heterogeneity, or even reflect an internal motility “clock” that alternates between stop and go states has been debated ^16,29^. The inability to disentangle various factors affecting motility *in vivo* has made it especially difficult to interpret T cell motility within crowds, as crowd dynamics are well-known to depend strongly on environmental structures ^65^ and mechanisms of agent-agent interaction ^66^, on top of agent-intrinsic behavior. Intriguingly, here we found that even in a homogeneous environment in the absence of environmental obstacles and without touching other cells, T cell motion is heterogeneous. While this heterogeneity at the single-cell level could be expected to impede motility in a crowd as faster cells “queue up” behind slower ones, experiments instead revealed that individual T cells synchronized to form stable, motile trains moving at speeds almost equivalent to those of cells moving alone. We showed that the ability of highly heterogeneous individuals to form stable, motile trains requires not only stabilization of cell-cell contacts through adhesion, but also a form of cooperation by which faster cells facilitate forward movement of slower ones. Together, the emergent crowd behavior we observed in single-lane traffic is surprisingly consistent with flocking theory as proposed by Reynolds ^42^ and Vicsek ^43^, which posit that flocking motion can be generated by agents that are cohesive and match their velocities. The model by Reynolds ^42^ also contains a third “separation” force that avoids making contacts between agents, which is required for fluid motion of animal flocks but is dispensable for flocks of T cells.

Our results are relevant to understanding T cell motility *in vivo*. Several studies have reported T cells navigating channel-like structures, such as in liver sinusoids ^67^ and in the recently discovered lymphoid structures in zebrafish where T cells navigate a tessellated network at high densities ^68^. In such settings, we would expect crowded motility to depend on cooperation. Moreover, as shown by the starkly different behavior of neutrophils, and corroborated by our *in silico* simulations, maintenance of motility is not a foregone conclusion of the geometry of our channels nor a general property shared by all motile leukocytes. Our data also raise the intriguing possibility that T cells may benefit from the presence of other T cells to move efficiently even if their synchronization is less overt than in channels. For example, T cells in lymph nodes and other tissues may require the ability to push other cells to limit crowding effects in mixed populations of slower and faster cells ^6^. Indeed, naïve T cells in lymph nodes have been found to subtly synchronize with very close-by neighbors, although force transmission or cell heterogeneity have not been investigated in this context ^16^.

Interestingly, the differences in movement synchronization between T cells and neutrophils may underlie their distinct functions. T cells have to keep moving in settings where they are crowded. By contrast, neutrophils are mostly found in crowds when they swarm to sites of tissue injury or infection where, upon arrival, they generally slow down substantially to contribute to the local barrier and repair response ^52,53^. It will be interesting to determine the underlying molecular mechanisms that drive continued motion in aggregating T cells, or those that lead to the stopping behavior in neutrophils. One possibility is that touching T cells synchronize waves of polymerizing actin as has been shown in cultured adherent endothelial cells ^69^. T cells and neutrophils may also differ in how they transmit forces resulting from direct cell-cell contact. In neutrophils, a mechanism similar to contact-dependent inhibition of locomotion might explain reduced motility and stability of cell clusters ^70,71^, for instance through triggering a mechanically-induced calcium flux, while T cells may respond differently to mechanical input. A better understanding of possible receptor-ligand signals triggered by cell-cell interactions between crowded neutrophils or T cells, as well as the likely differences in the impact of mechanical input, is an important area of future investigation.

Together, our work adds a new perspective to the field of T cell migration by presenting evidence that T cells cooperate to move effectively in tight, crowded spaces. To date, *in silico* models of T cell migration either did not explicitly account for cellular crowding or simulated cells at low density ^5,58,59,72–76^, or specifically enforced motility in dense crowds ^77,78^. Through the lens of crowd dynamics, dense T cell collectives appear to be a remarkably resilient active matter system and could provide an interesting case study for efficient crowd motility. The fundamental diagram places T cells among other social crowds such as ants, even though T cells differ greatly from these systems in that they are deformable, highly heterogeneous, and are able to interact through direct physical contact. Interestingly, the large differences in cell-cell interaction between T cells and neutrophils were mostly obscured in the fundamental diagram focusing only on distance to the nearest neighbor. Explicitly considering clusters of agents in direct physical contact may therefore reveal further differences not only between various motile cell types, but also between other active-matter systems. As T cells encounter crowds in many of the environments they traverse, further determining the mechanisms of T cell crowd control will be crucial to our understanding of T cell migration in health and disease.

## Data and code availability

Data and code required to reproduce the analyses in this manuscript will be made available on Github and Zenodo on publication. In the meantime, resources are available on reasonable request to the corresponding authors.

## Author contributions

JNM, DRP and JT conceptualized the research and obtained funding. JP and JNM designed the experimental system. Experiments were performed by JP, MH, MaM, AB, and AC. MiM processed the video data and annotated cell tracks and contacts with help by IMNW, JT and LEW. IMNW, DRP and JT performed formal analyses on experimental data. IMNW and JT designed and implemented the computational models; LiW fitted models to experimental data supervised by IMNW. IMNW and JT analyzed model outcomes. IMNW, JP, DRP, JNM and JT designed data visualizations. IMNW, JNM and JT each drafted and edited parts of the manuscript, and all authors reviewed the final manuscript.

## Acknowledgements

We gratefully acknowledge the Flow Cytometry Core Facility (FCCF) and Advanced Bioimaging Facility (ABIF) of the Life Science Complex at McGill University. We thank Karel Prud’homme and the staff from the Comparative Medicine and Animal Resources Centre (CMARC) for their care of our mouse colony at McGill University. We are also grateful to Dr. Heather Melichar (McGill) for providing sectioned thymus tissue for analyses and her helpful feedback, along with all Mandl lab members for their input.

## Materials and Methods

### Ethics

#### Human tissue samples

In this study, we re-analyzed immunohistochemistry images of human primary melanoma and tonsil samples that were obtained and imaged in a previous study ^79^, in accordance with the regulations of the Dutch Committee on Regulation of Health Research.

#### Mice

Nucleus-signal tagged TdTomato fluorescent reporter mice, nTnG (JAX number: 023537) ^80^ were obtained from The Jackson Laboratory and crossed to F-actin fluorescent reporter mice, Lifeact-GFP ^81^, that were generously shared by Janis Burkhardt (University of Pennsylvania), both on a C57Bl/6 background. C57Bl/6 were obtained from The Jackson Laboratory (JAX number: 000664). All mice were bred in-house under SPF conditions in the animal facility at McGill University at a temperature of 18-24°C, 30-70% humidity, and 12h/12h light-dark cycles. Mice were used for experiments at 8-12 weeks of age. Both male and female mice were used. All mouse studies were performed in accordance with the Guide for the Care and Use of Laboratory Animals, with approval by the McGill University Facility Animal Care Committee (protocol number MCGL-7570).

### Immunohistochemistry and quantification of T cell density in tissues

Inguinal lymph nodes and thymus isolated from C57Bl/6 mice were fixed in buffer containing 0.05 M phosphate buffer, 0.1 M l-lysine, 2 mg/mL NaIO4, and 10 mg/mL paraformaldehyde overnight, followed by 30% sucrose. Tissues were frozen at -80°C in Tissue-Tek OCT compound, and were later sectioned into 10-20 μm thick slices using a Leica CM3050 S cryostat. The sections were blocked with buffer containing 0.1 M Tris, 1% FBS, 0.3% Triton X-100, and 1% gelatin from cold water fish scales. Lymph nodes were stained overnight with a primary antibody against ERTR7 (Novus Biologicals), followed by a Donkey anti Rat AF647 secondary antibody (Jax) and a CD3-AF594 conjugated antibody (BioLegend). Lymph node sections were mounted with Fluoromount-G Mounting Medium with DAPI (Invitrogen). Thymic lobes were stained with a CD4-AF647 conjugated antibody (BioLegend), a primary CD8 antibody (BioLegend) followed by a goat anti-rat AF555 antibody (Invitrogen) and a primary Keratin-5 antibody followed by a goat anti-rabbit AF488 (LifeTech). Thymic sections were stained with DAPI (Sigma) and mounted with ProLong Gold Antifade Reagent (Invitrogen). Images were acquired with a Zeiss LSM880 confocal microscope using a Plan-Apochromat 40x (NA = 1.3) oil objective.

For all datasets, we selected areas of high cell density (human samples: 3 regions per specimen; mouse samples: two regions per specimen; 3 biological replicates per tissue type). For each sample, we determined T cell positions in a squared observation window of 75×75 µm. The cell positions were detected using ImmuNet ^79^. The distance to the nearest neighbor was estimated using the function *Gest* (R package *spatstat* v3.0.7^82^), which corrects for the size and shape of the observation window.

### Migration assays in micro-fabricated channels *in vitro*

#### Microchannel fabrication

Polydimethylsiloxane (PDMS) RTV-615 (Newark Electronics, #00Z716) was mixed with curing agent (ratio 10:1), degassed by centrifugation at 1000 g for 1 min, poured into custom epoxy molds manufactured by 4DCell, and further degassed under negative pressure within a vacuum desiccator for at least 1 hour. After 2 h curing at 80°C, PDMS chips were carefully removed from molds using isopropanol, blow-dried using a gas duster, punched to open the channels (Harris Uni-Core, #WHAWB100029), dusted with clear tape, and further cleaned in an ultrasonic bath (VWR, #14003-062) filled with pure ethanol for 3 min. Chips were then blow-dried and functionalized with air plasma for 2 min (Harrick Plasma, #PDC-001) to be bonded to functionalized glass-bottom dishes (WPI, #FD35-100). The resulting devices were baked using an oven for 1 hour at 60°C. Microchannels were functionalized in air plasma for 2 minutes and coated with fibronectin (Sigma-Aldrich, #F-1141, 10 µg/mL in PBS), for 1 h at 37°C, 5% CO_2_. Before use, all devices were washed twice in PBS and twice in complete media w/o phenol red, and incubated in complete media w/o phenol red for at least 30 min at 37°C, 5% CO_2_.

#### T cell migration assays

Lymph nodes from Lifeact-GFP nTnG mice were harvested and crushed through a 70 µm filter in StemCell buffer (PBS w/o Ca^2+^/Mg^2+^ supplemented with 2% FBS and 1 mM EDTA). T cells were isolated in FACS tubes using the EasySep Mouse T Cell Isolation Kit (StemCell, #19851) following the manufacturer’s instructions with an additional step of separation over the magnet. After enrichment, T cells were washed in complete media (RPMI 1640 supplemented with 10% heat-inactivated fetal bovine serum (FBS), 100 U/mL penicillin, 100 µg/mL streptomycin, 2 mM L-glutamine, 1 mM sodium pyruvate, 10 mM HEPES, 1X non-essential amino acids, and 5 µM 2-mercaptoethanol), adjusted to 2.5×10^6^ cells/mL, and supplemented with 2 µg/mL αCD28 (Biolegend, #102116). The cell suspension was distributed over a 96-well plate (200 µL/well) that had been precoated with 3 µg/mL anti-CD3ε (Biolegend, #1000340) for 2 h, 37°C, 5% CO_2_. Two days later, the cells were harvested, washed, resuspended in complete media supplemented with 20 ng/mL rm-IL-2 (Biolegend, #575406), and cultured in a T75 flask at 5×10^5^ cells/mL. Activated T cells were used two days later, at day 4 post activation. Microchannel access ports were loaded with 10 µL of activated T cells resuspended at 10^8^ cells/mL in complete media w/o phenol red on one side and 10 µL complete media w/o phenol red on the other. No chemokines were added to the microchannels. Before imaging, loaded devices were incubated at least one hour at 37°C, 5% CO_2_.

#### Neutrophil migration assays

Bone marrow from Lifeact-GFP nTnG mice was harvested by flushing femurs and tibia bones with complete media and passed through a 70 µm filter. Red blood cells were lysed by resuspending the bone marrow in 1 mL of ACK Lysing Buffer (Thermo Fisher Scientific, #A1049201) for 3 minutes. Neutrophils were isolated using the EasySep Mouse Neutrophil Enrichment Kit (StemCell, #19762) following the manufacturer’s instructions. Neutrophils were resuspended at 1×10^7^ cells/mL in complete media w/o phenol red supplemented with 20 ng/ml GM-CSF and then pulsed with 1 µg/ml LPS (InvivoGen, # tlrl-smlps) for 30 minutes. After washing, the cells were resuspended in complete media w/o phenol red supplemented with 20 ng/ml GM-CSF at 2.5×10^8^ cells/mL. Microchannels were loaded as described for T cells.

#### Image acquisition

Cell migration was recorded by time-lapse widefield microscopy using an Axiovert 200M Fully Automated Inverted Microscope (Zeiss) equipped with a 20X/0.8 NA Plan-Apochromat objective (Zeiss), a top-stage incubation system set at 37°C and delivering 5% CO_2_ in humidified air (Live Cell Instrument, #Chamlide TC-L-Z003), and a monochrome camera (Hamamatsu). Fluorescence emitted from GFP and tdTomato was collected from a single z-stack using an X-Cite 120 LED (Excelitas Technologies) combined with Zeiss filter sets (GFP: FS10, tdTomato: FS15). Images were acquired every 20 seconds for 30 minutes to more than 1 hour. Movies were acquired using ZEN (Zeiss).

#### Annotation and tracking

Cells were detected using StarDist ^83–85^ and tracks were generated manually using in-house software. Contacts between cells (i.e., cells were touching each other as visible from the Lifeact-GFP signal) were also manually annotated for each frame.

### Analysis

#### Track preprocessing

To compensate for rotation differences, tracks from each movie were rotated to align the average direction of movement with the x-axis for 1D analysis. To remove noise in cell positions, tracks were smoothed with a Kalman filter (R package *dlm* v1.1.6^86^; using function *dlmSmooth* with a second order polynomial model, *m*_0_ = [0, 0], *C*_0_ = [[0, 0], [0, 10]], *dV* = 5, *dW* = [0, 2]).

To factor out variability between biological replicates, tracks from each movie were rescaled before pooling by multiplying positions with a constant factor:

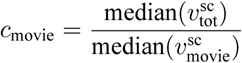

with medians over the single-cell (sc)-track speeds within one movie and across all movies, respectively.

#### Fundamental diagram

For the fundamental diagram of ants, densities and speeds were extracted from published data ^37^ using a plot digitizer. For the fundamental diagrams of cars ^38^ and pedestrians ^87^, raw tracks were available and processed as follows: speeds were computed as 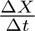, with Δ*X* the agent displacement between consecutive recordings Δ*t* time apart. Local densities were computed as 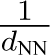, with *d*_NN_ the distance to the nearest neighbor agent. Speeds and densities for T cells were computed the same way (for densities, only the distance to the nearest neighbor from the same channel was considered). Fundamental diagrams were then obtained as the rolling average of speeds for increasing densities (window size *k* = 3000 for all data except ants, where *k* = 20 was used since there were much fewer data points available). To obtain a normalized version of the fundamental diagram, speeds were normalized relative to the maximum of the observed rolling averages. Densities were multiplied with the characteristic agent length *ℓ*_0_ so as to achieve a density of 1 when *d*_NN_ = *ℓ*_0_. For *ℓ*_0_, the following values were used: 5 m for cars (averaged from the MiTra dataset ^39^), 1.8 mm for ants (as reported in John et al ^37^), 0.25 m for pedestrians (estimate), and 12 and 16.9 µm respectively for neutrophils and T cells (the median distance between contacting cells).

#### Single-cell sc-track extraction

To analyze motility of single, non-touching cells, single-cell sc-tracks were extracted for all intervals where cells moved independently for at least 15 frames (5min); potentially yielding multiple sc-tracks per cell.

#### Survival analysis of persistent intervals

Because many cells traversed the channels without ever turning, persistence time was analyzed by treating it as the “survival time” of a given movement direction, thus accounting for intervals where the end of this persistent state was unobserved (e.g. because the video ended, cells left the field of view, or sc-tracks touched other cells and were thus no longer sc-tracks). For each track, the direction between each pair of consecutive frames was extracted by considering the sign of the (Kalman-smoothed) instantaneous velocities. These directions were then split into intervals where cells maintained the same direction, which were either “censored” (at the end of the sc-track) or fully observed. Unbiased estimates of the persistence time were then obtained using the Kaplan-Meier estimator (R package *survival* v.3.5.3^88,89^) and visualized in a Kaplan-Meier curve (R package *ggfortify* v.0.4.16^90^).

#### Motility analysis of sc-tracks

To assess speed heterogeneity within and between sc-tracks, instantaneous “step” speeds were first computed within each sc-track (R package *celltrackR*, v.1.2.0^91^) to obtain the standard deviation and average of speeds per sc-track. The arrest coefficient was computed as the fraction of instantaneous speeds that were below 5 µm/min, as this threshold gave good correspondence of the inferred “stop” state with those observed when looking at the video. Mean squared displacement (MSD) curves for all sc-tracks were computed using celltrackR and fitted using Fürth’s equation in 1D to obtain single-cell estimates of the motility coefficient and persistence time ^92,93^.

#### Long-distance interactions of single cells

To investigate potential long-range communication between cells in channels, two analyses were performed. First, the fundamental diagram (see above), showing instantaneous speed as a function of (inverse) distance to the closest neighbor cell was computed. If cells actively adjust their speed based on long-range communication, this should result in a distance-dependent trend in the fundamental diagram. Second, to investigate the possibility that cells actively change their direction in response to long-range communication, each sc-track was analyzed to determine whether its direction was aligned with that of the closest neighboring cell or train, reporting the percentage alignment as a function of nearest-neighbor distance.

#### Train sizes

To extract cell “trains”, a graph was constructed for each video frame where cell positions were nodes, and edges were added between contacting cells according to the manually annotated contact information. Trains were then extracted as the connected components within these graphs (R package *igraph* v.1.5.0^94^).

#### Analysis of train motility

To analyze motility of trains, track segments were extracted where cells were in trains of a given composition as described for sc-tracks above. Instantaneous velocities of a given train were then measured as the average of the instantaneous velocities of all cells in the train. Average train velocities were then computed as the average of the train instantaneous velocities over all steps where the train existed in that same composition, and arrest coefficients were calculated in the same way as for sc-tracks. Speeds of trains of length *N* were compared to a “simple queueing model” in which *N* cell speeds were drawn from the sc-track speed distribution and the minimum was taken – representing a queue of cells moving in the same direction and unable to push each other, where the slowest cell would be in the front.

#### Train survival analysis

Similar to the persistence time of sc-tracks, the “survival time” of trains was difficult to measure since many trains left the field of view before decaying. Analogous to the analysis of persistence time described above, we therefore assessed train stability by constructing Kaplan-Meier curves of train survival, censoring intervals at the beginning and end of the movie, or when one or more of the train’s cells entered or left the field of view. In addition, since the growth of trains is highly dependent on the number of cells in a given movie or simulation, we focused on train decay only, censoring intervals also when other cells joined the train such that the survival time of the original train could no longer be observed.

#### Cell collision analysis

Collision events were defined by searching for two cells that met the following criteria: both cells were fully visible and not in contact with each other, for five minutes or longer before the event. Cell-cell distance was measured as the absolute difference in the x-position between the two cells; cell pair speed was computed as the average of the central finite differences of each cell’s x-position, averaged across both cells. From these collision events, a collision was classified as head-to-tail if (a) both moved in the same direction before contact; (b) the rear cell (s1) was strictly faster than the front cell (s2) at every timepoint in the 2 minutes before contact.

### Cellular Potts Model simulations

#### Overview

Cellular Potts Models (CPMs) have been described in detail elsewhere ^45,46^. Briefly, CPMs describe cells in terms of occupied pixels *p* on a grid by assigning each pixel *p* the “identity” *σ*(*p*) of the cell to which it belongs (with *σ*(*p*) = 0 for unoccupied pixels). Cells move through *copy attempts*, which occur sequentially from a randomly sampled “source” pixel *p_s_* to a “target” pixel *p_t_* sampled from the Moore neighborhood of *p_s_*. Thus, a cell *i* can either “protrude” (*σ*(*p_s_*) = *i, σ*(*p_t_*)≠*i*), “retract” (*σ*(*p_s_*) = 0*, σ*(*p_t_*) = *i*), or be “displaced” by another cell *j* (*σ*(*p_s_*) = *j, σ*(*p_t_*) = *i*). The balance between those options is governed by the success probability:

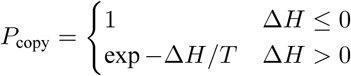

which depends on the *temperature T* (a parameter controlling the amount of stochasticity) and the change in system *energy H* that would occur if the copy attempt were to succeed. “Favorable” copy attempts that lower the energy are favored (*P*_copy_ = 1), while “unfavorable” changes have a lower success rate.

What is “favorable” or “unfavorable” is then dictated by the choice of the *Hamiltonian* energy *H* which is set by the modeller. The Act-CPM ^47,48^ starts from a simple model of cell biophysics based on surface tension along with area and perimeter preservation ^45,95^. To model active cell migration, the Act-CPM extends this baseline model by considering symmetry breaking due to polarized actin polymerization activity within the cell. Whenever a pixel *p* is assigned to a new cell σ ≠ 0, it gets an “actin polymerization activity” *A*(*p, t*^∗^) = max_act_, which then decreases over max_act_ steps:

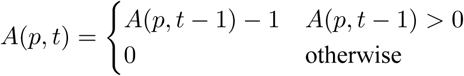

While pixels retain their actin polymerization activity, they are more likely to succeed at protrusive copy attempts because of an additional contribution Δ*H*_act_(*p_s_* → *p_t_*) that favors copy attempt from “more active” sources *p_s_* into “less active” target *p_t_*.

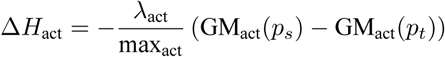

Here, GM_act_(*p*) is the geometric mean of the local “protrusive activity” *A*(*p*) at pixel *p* and its (Moore) neighbor pixels belonging to the same cell. Thus, this term is negative (i.e., energetically favorable) whenever

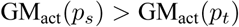

Note that although CPMs are defined by the system energy *H*, the difference in energy Δ*H* caused by a potential copy attempt can be interpreted as a locally applied force ^46^. Δ*H*_act_ can be loosely interpreted as the force exerted by polymerizing actin on the membrane at the front of the cell. The parameter *λ*_act_ controls the relative magnitude of this force compared to the other forces acting on the cell (e.g. those arising from surface tension). The other migration parameter, max_act_, determines the stability of the polarized state.

#### Model extensions

In this work, we extend the Act-CPM with one additional parameter *η* that represents the efficiency at which a protrusion force from one cell is transferred to another:

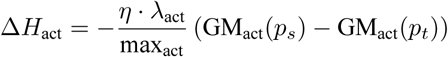

where:

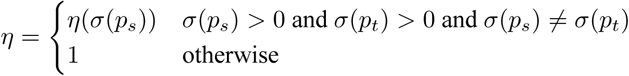

*η*(*σ*) is a parameter with a value between 0 and 1, which only affects protrusive copy attempts from one cell into another cell. When *η* = 1, the model is equivalent to the Act-CPM. Lower values of *η* effectively scale down the “effective force” that can be exerted on another cell by the protrusion of cell *σ*(*p_s_*). In the most extreme case (*η* = 0), cell *σ*(*p_s_*) cannot transmit any of the actin-generated force to cell *σ*(*p_t_*), yielding a scenario with no cooperativity since *σ*(*p_t_*) is unaffected by the movement of cell *σ*(*p_s_*).

#### Adhesion and contact stabilization

Cell-cell contacts in the CPM can be easily stabilized through the adhesion term in the Hamiltonian ^45^, which assigns an energetic penalty to every pair of neighboring pixels that do not belong to the same cell. The size of this penalty depends on the type of cells in contact; we distinguish *J*_cell,cell_ for contacts between cells and *J*_cell,bg_ for contacts between the cell boundary and empty background. Here, we use *J*_cell,bg_ = 0 (see **Table S4**). Unless otherwise specified, we set *J*_cell,cell_ = 0 to the same value such that cell-cell contacts are energetically neutral compared to cell-background contacts. For simulations with adhesion parameter *γ*, we instead set *J*_cell,cell_ = −*γ*, which favors and stabilizes cell-cell contacts.

#### Parameter selection

Most model parameters were selected based on published work (see **Table S4**). To fit motility of single cells in the CPM to the motility of sc-tracks, smoothed and normalized sc-tracks from all videos were pooled. To match not only the average but also the variation in motility across sc-tracks, each cell’s variable parameters *c_i_* = (*λ*_act_, max_act_)*_i_* were sampled from a bivariate lognormal distribution:

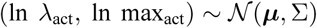

with means ***µ*** = (*µ_λ_, µ_m_*) and a covariance matrix Σ with diagonal elements (*σ_λ_, σ_m_*) and covariance *σ_λm_*. Thus, a heterogeneous cell population is defined by five parameters: ***θ*** = (*µ_λ_, µ_m_, σ_λ_, σ_m_, σ_λm_*). To fit ***θ*** to the observed distributions of speeds and arrest coefficients of sc-tracks, Approximate Bayesian Computation (ABC) based on sequential Monte Carlo (SMC) as described by Sisson et al. ^96^ was used. We outline our full implementation in the Supplementary Materials, along with the choice in prior and distance function.

### Simulations of head-to-head and head-to-tail collisions

These CPM simulations were performed using a custom-written parallelized implementation of the CPM written in JavaScript and WebGPU. Simulations were constructed at a resolution of 0.454 µm/pixel and 0.5 second/MCS. Microchannel “walls” were placed 13 pixels (6 µm) apart and were implemented as “empty” background pixels of a special type, where copy attempts *into* the channel walls are forbidden (*P*_copy_ = 0) whereas copy attempts *from* the channel walls proceed as usual. A burn-in phase of 4500 MCS was used.

For head-to-head CPM simulations, channel length was set to 512 pixels and non-wrapping boundary conditions were used. Each channel was initialized with two cells: one placed at x = 20 pixels (≈9 µm from the left wall) and one at x=494 pixels (≈9 µm from the right wall). This placement forces the cells to develop polarity towards the middle, and eventually collide. Both cells were assigned the same parameters (Supplementary Materials and max_act_ = 45*, λ*_act_ = 450).

For head-to-tail CPM simulations, each channel was initialized with two cells moving right. The faster “rear” was seeded at x=20 pixels and used default parameters, including max_act_ = 45 and *λ*_act_ = 450. The slower “front” cell was seeded at x=92 pixels and used a lower value of *λ*_act_ = 161. A burn-in phase of 1200 MCS was used. This setup does not yet guarantee that a head-to-tail collision actually does occur, since the slower cell could move towards the left. Therefore, the trajectories of both cells were smoothed with splines and differentiated to estimate cell speeds and then classified as follows: a simulation run was considered a head-to-tail collision if, at every pre-collision timepoint, the left cell moved towards the right and the left cell was faster than the right cell. Further, the right cell had to be moving towards the right at the frame immediately before contact. Random seeds where this occurred were saved and simulations at different values of *η* and *γ* (which only affect the post-collision dynamics) were run for these saved seeds.

### Detailed simulations of T cell motion in microchannels

These CPM simulations were performed in Artistoo ^97^. Simulations were constructed at a resolution of 0.454 µm/pixel and 0.5 second/MCS, i.e., as in the collisions simulations. Microchannel walls were implemented the same way as in the collision simulations. Simulations were matched to the corresponding movie in terms of size of the imaging window, size and placement of microchannels, and total duration.

At the beginning of the simulation, cells were placed in the same positions as the cells in the first frame of the corresponding recorded movie. Cells were seeded as a single pixel. After a burn-in time of 50 simulation steps (where only copy attempts affecting the current cell are allowed to let this cell attain its desired area), we gave each cell an initial direction matching the initial direction of the corresponding T cell. This was done by setting all pixel activities *A* to zero on either the left side of the cell (causing initial movement the right) or on the right side of the cell (causing initial movement to the left).

The simulation was given open boundary conditions by implementing a 100-pixel “buffer zone” both on the left and right of the simulated field of view. Cells were removed from the simulation once all their pixels were inside the buffer zone on either side (and thus, outside of the “imaging field of view”). After each simulation step, new cells were seeded if the total number of cells remaining in the simulation was below the number of cells recorded in the corresponding movie at that point in time. To seed a new cell, a channel and “entry side” (left or right) was chosen randomly, and a new cell was initialized in the buffer zone with initial direction towards the field of view as described above (if the selected buffer zone position already contained a cell, a different location was sampled). This set-up allowed us to match the overall cell density in the simulation, while still letting outcomes emerge stochastically (since the site where cells leave the channels emerges from CPM update dynamics and the new entry sites are chosen randomly).

Every 20 seconds (40 MCS, matching the frame rate of movie recordings), the centroids of all cells within the field of view were recorded, as well as all pairs of cells that were “in contact” (i.e., cell *σ*_1_ and cell *σ*_2_ are in contact iff there exists at least one pair of (Moore) neighbor pixels (*p_i_, p_j_*) for which *σ_i_* = *σ*_1_ and *σ_j_* = *σ*_2_). The resulting tracks and annotated contacts were then analyzed in the same manner as those extracted from the experimental data.

