## Supplementary Methods, Figures and Tables for "Cooperative motility emerges in crowds of T cells and prevents jamming"

### Supplementary Material

June 26, 2026

### Supplementary Figures

#### Figure S1: Replicate movies are variable in cell density and motility.

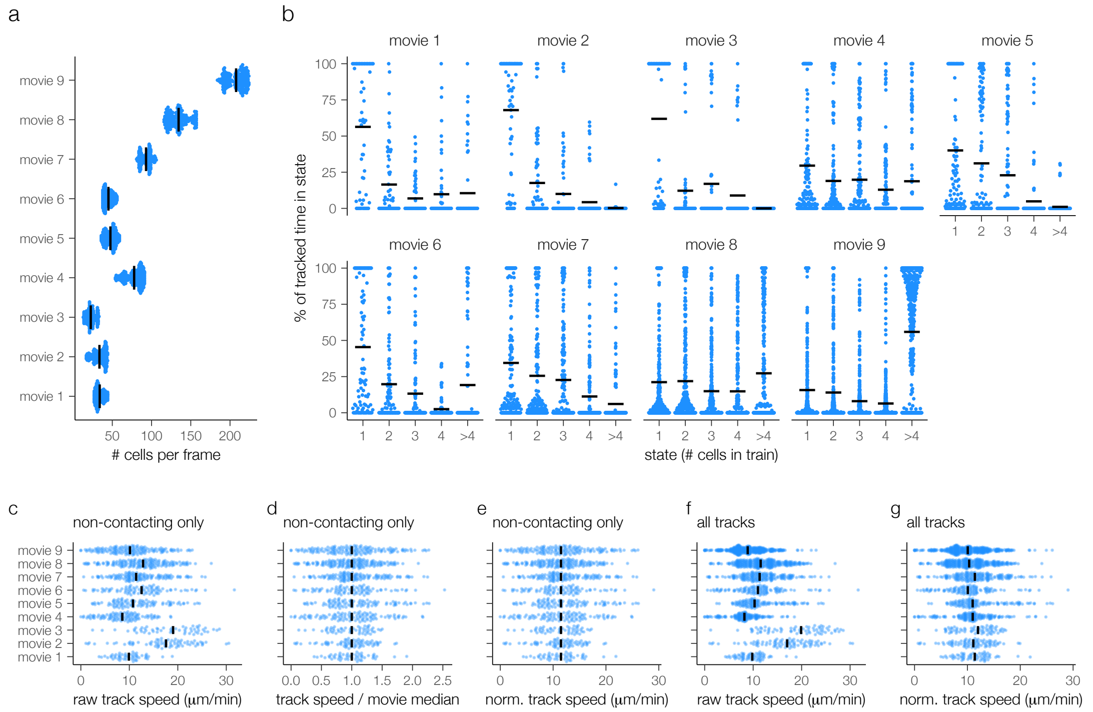

**(a)** Movies were variable in terms of cell densities (points: number of cells for each frame, lines: movie average of cells per frame). **(b)** Variable densities translated to a variable distribution across train sizes. Each dot is one track, showing the percentage of time spent in each state. Horizontal black lines: average across all tracks. **(c)** Also motility was variable between biological replicates, with different speed distributions even for single-cells that did not touch other cells. **(d,e)** To focus on density-dependent variability rather than biological variability, we computed a scaling factor for each movie to equalize the sc-track median to the overall median (Methods). **(f,g)** All track speeds in the dataset were multiplied with this scaling factor to obtain the rescaled dataset used for further analysis.

**Figure S2: Fundamental diagrams of T cells compared to different systems.**

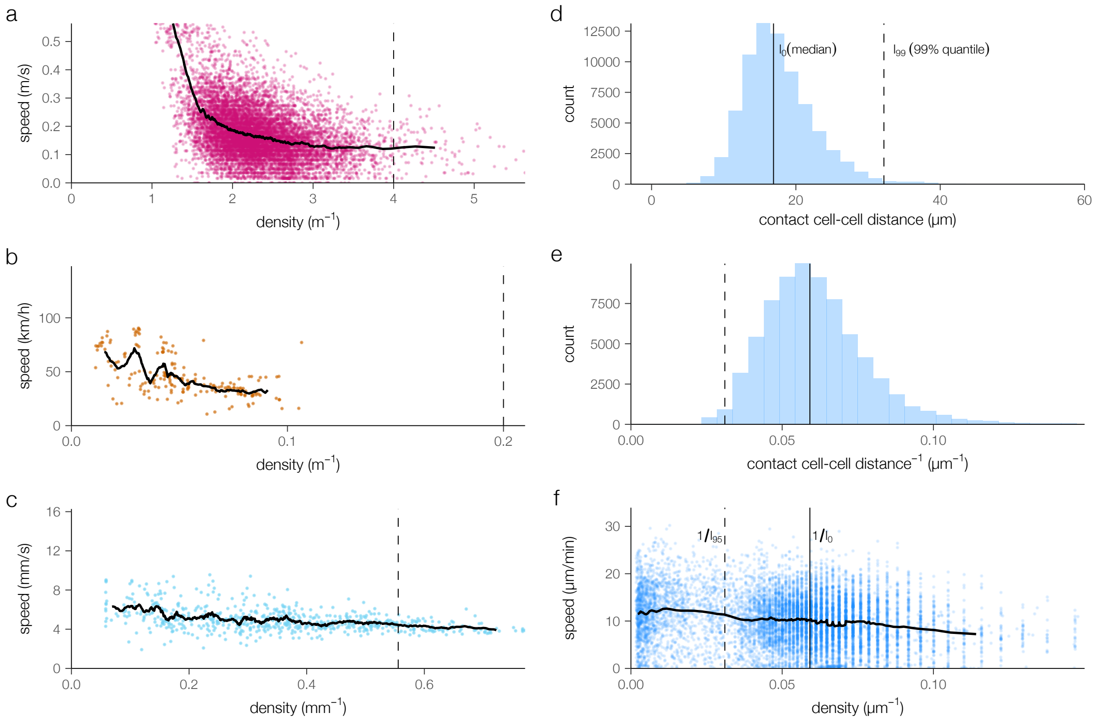

**(a-c)** Fundamental diagrams of pedestrians (a), cars (b), and ants (c) as in Fig. 1, now showing the underlying point distribution before computing the moving average (lines). Maximum 10000 points are shown to prevent overplotting. **(d,e)** Distribution of distances between pairs of cells in direct contact (d) along with the corresponding densities (e). Dashed vertical line indicates the density where cells start touching. **(f)** Fundamental diagram of T cells showing the underlying distribution (a random sample of 10000 points from the data is shown).

**Figure S3: An extended queuing model of single-lane T cell traffic explains single-cell, but not train dynamics.**

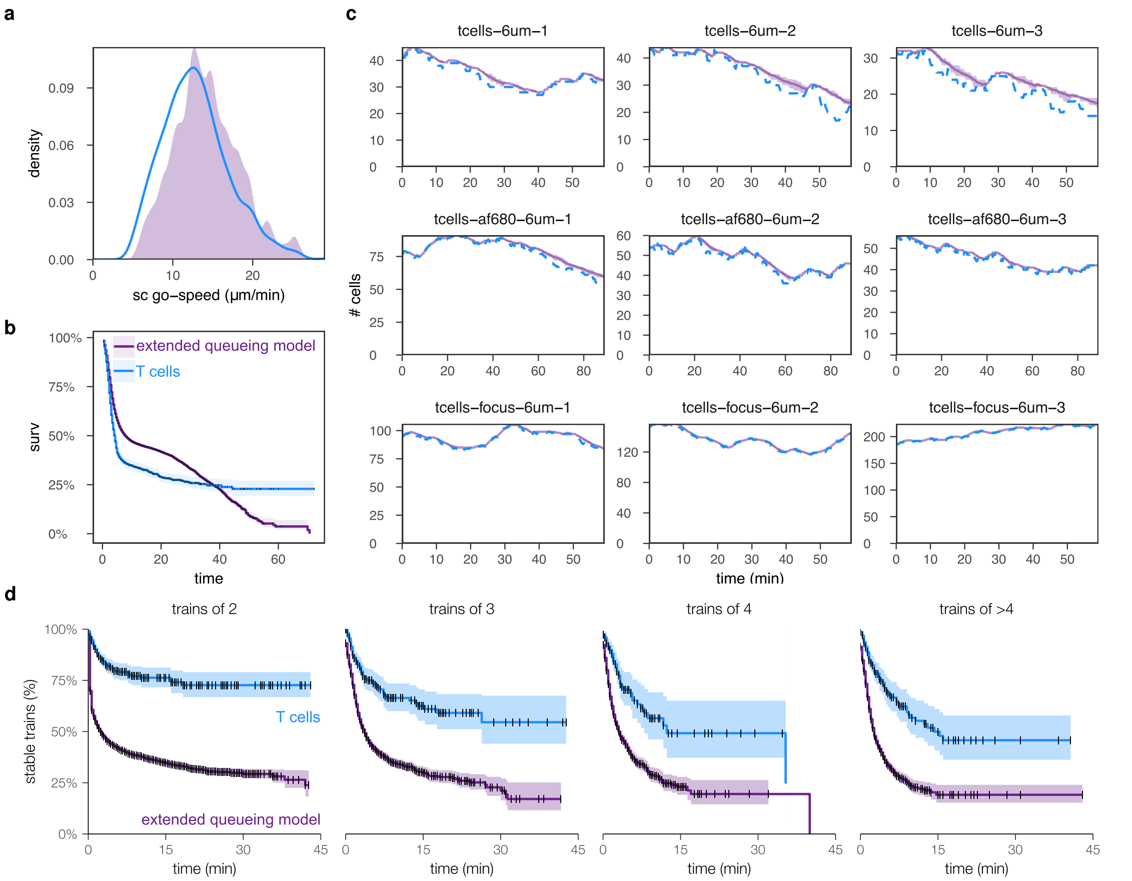

**(a,b)** The sc-track distribution (a) and survival of directionality (b) compared between T cells and the extended queuing model fitted on sc-tracks (20 simulation twins per replicate movie; total 180 simulations). **(c)** Cell entry in the model was set to match the number of cells in each movie over time as closely as possible. **(d)** Survival of trains of various sizes compared between experimental data and extended queuing model.

**Figure S4: No evidence for long-range interactions.**

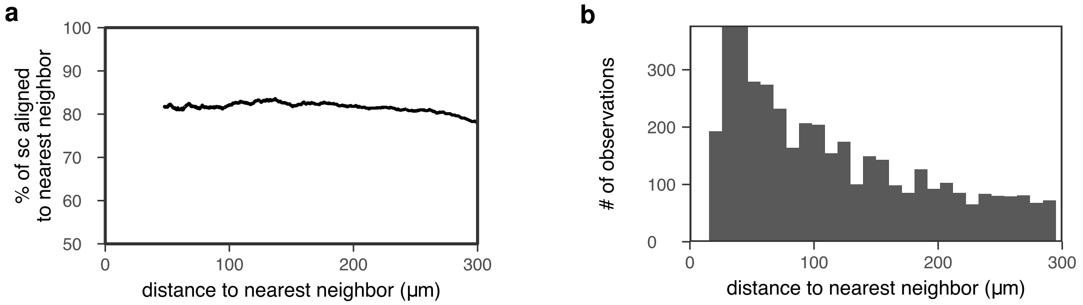

**(a)** To assess potential long-range interactions beyond the fundamental diagram, we examined directional alignment between single cells and their nearest, non-touching neighbor in the same channel (which may or may not be a single cell). For each such case, we recorded whether or not the directions were aligned (yes/no). The graph shows the percentage of aligned cases as a function of nearest-neighbor distance, computed via a rolling average over 1000 cases. Although the alignment is generally high, indicating global directional bias, there is no evidence of long-range sensing as there is no distance-dependence of the alignment. **(b)** Shows for each distance the number of cases observed, based on which (a) was computed. Data in this figure are based on pooled normalized tracks from all 9 replicate movies.

#### Figure S5: Fitting the Act-CPM to sc-motility.

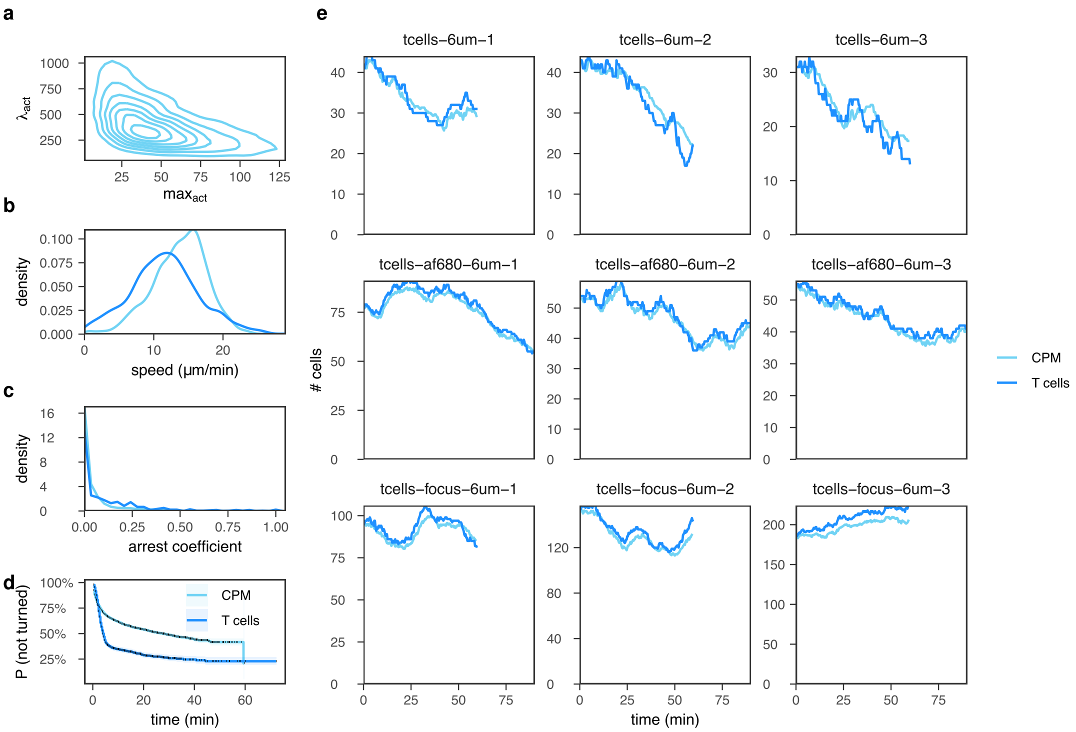

**(a)** To accomplish heterogeneous motility in the CPM, its main motility parameters $max_{act}$ and $\lambda_{act}$ were drawn from a bivariate lognormal distribution that was fitted to match motility of pooled, normalized sc-tracks from all 9 replicate movies. **(b-d)** sc-tracks from the CPM with parameters drawn from the fitted distribution indeed roughly match T-cell sc-tracks in terms of their heterogeneous speed distribution (b), arrest coefficients (c), and persistence (d). CPM data are from 3 independent simulation twins per replicate movie (i.e., 27 simulations in total). **(e)** As with the extended queuing model (Fig. S3), cell numbers were matched to the movies over time.

#### Figure S6: Statistics and normalization of neutrophil tracks.

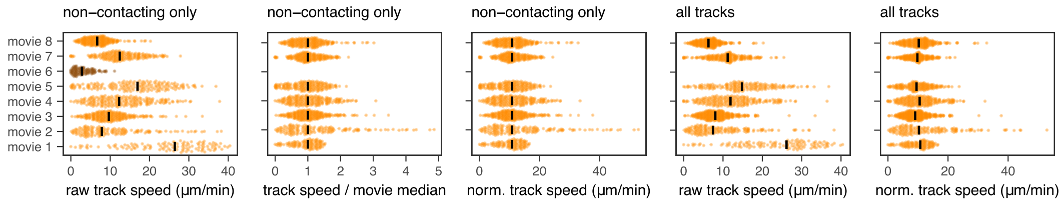

We recorded 8 movies of neutrophils in microchannels, which like T cells were variable in motility of single cells. Because there was one movie in which single cells barely moved at all (even single cells), that movie (movie 6, dark) was excluded from further analysis. Movies were rescaled as described previously for the T cell data (Fig. S1).

#### Figure S7: Neutrophil train slowdown appears to match a simple queuing model.

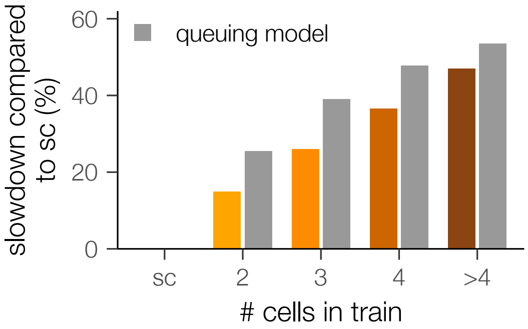

The same analysis as in Fig. 4e performed for the neutrophil data from Fig. 6d.

#### Figure S8: The fundamental diagram of neutrophil motion in single-lane traffic.

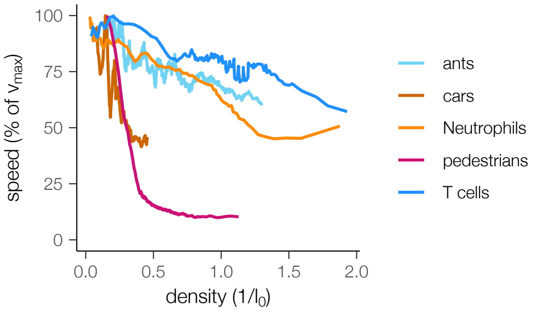

As Fig. 1i, now adding the speed-density relation for neutrophils (computed on pooled, normalized neutrophil sc-tracks from 7 movies as for T cells in Fig. 1i). Density normalization was now performed using the neutrophil median contact distance (12 μm). Since the fundamental diagram focuses on relations over longer distances, the difference between T cells and neutrophils is not nearly as clear here.

### Supplementary Methods

#### Extended queuing model simulations

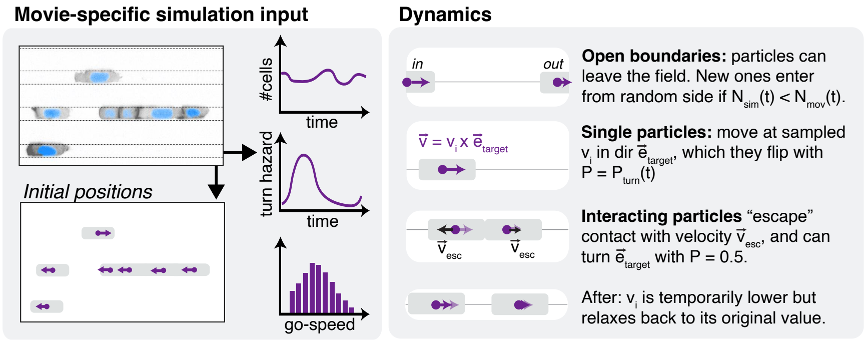

Figure A1: Model overview.

**Model overview.** Our "extended queuing model" of T cell motility in crowded microchannels under minimal cell-cell interactions is based on the contractile particle model by Baglietto and Parisi [1]. An overview of model parameters is given in **Table S3** and an overview of the model in **Figure A1**.

In the Baglietto model, single particles move at a desired velocity $\vec{v}_i$ with magnitude $v_i(t)$ and direction $\vec{e}_{\text{target}}$. They also have a dynamic interaction radius $r_i(t)$ which is bounded between $r_{\text{min}}$ and $r_{\text{max}}$, and is related to movement speed as follows:

$$v_i(t) = v_i^{\text{max}} \left[ \frac{r_i(t) - r_{\text{min}}}{r_{\text{max}} - r_{\text{min}}} \right]^{\beta}$$

This mechanism ensures that $v_i(t)$ changes between 0 (when $r_i(t) = r_{\text{min}}$) and $v_i^{\text{max}}$ (when $r_i(t) = r_{\text{max}}$). In principle, moving particles have $r_i(t) = r_{\text{max}}$ and thus $v_i(t) = v_i^{\text{max}}$. But whenever two particles $j$ and $k$ get within distance $d_{jk} < (r_j + r_k)$, physical exclusion kicks in, where:

1. The velocity $\vec{v}_i(t)$ is temporarily set equal to an escape velocity $\vec{v}_i^{\text{esc}}$ with magnitude $v_e$ and direction $\vec{e}_{\text{esc}}$ away from the other particle, until the interaction is resolved and $d_{jk} > (r_j + r_k)$. When a particle is in conflict with multiple neighbors, the escape direction $\vec{e}_{\text{esc}}$ is the sum of the individual escape directions. (For our 1D lines, this only happens when particles have a neighbor within conflict distance on both sides — in which case $\vec{e}_{\text{esc}}$ equals the null vector because both components cancel each other out.)

2. Once the interaction is resolved, the particle once again moves in direction $\vec{e}_{\text{target}}$ — but its radius $r_i$ is set to $r_{\text{min}}$ and only slowly relaxes back to $r_{\text{max}}$, increasing every timestep $\Delta t$ with an amount:

$$\Delta r = \frac{r_{\text{max}}}{\tau / \Delta t}$$

until it either reaches $r_{\text{max}}$ or collides again with another particle. Here, $\Delta t$ equals 1 second, small enough to avoid artefacts as described by Baglietto and Parisi [1].

These interaction rules ensure that after escaping the other particle, each particle must accelerate to attain its desired speed $v_i^{\text{max}}$. The speed at which this happens is controlled by parameters $\beta$ and $\tau$.

**Agent turning.** In the abovementioned model, the specific target direction $\vec{e}_{\text{target}}$ must be specified depending on the system to be simulated. For the case of T cells, we define the following heuristics for turning in microchannels:

1. Particles can attempt to turn if, over a "memory" period of $M$ steps, they have displaced less than 10% of their "desired" displacement ($v_i^{\text{max}} \cdot M \cdot \Delta t$). This means they lose their intrinsic polarity if they do not succeed at moving for $M$ steps. The directional change occurs with probability $p_{\text{switch}}$ and succeeds only if there is no other particle within conflict distance in that direction.

2. Particles can also attempt to flip their target direction spontaneously. This again only succeeds if there is not another particle within conflict distance in that direction, and happens with a probability $p_{\text{turn}}(t_p)$ that depends on the current duration $t_p$ of the persistent interval (see *Parameter selection* below).

**Parameter selection.** Where possible, model parameters were based on observations in the T cell data. For example, to match the heterogeneous motility observed in T cells, each particle's $v_i^{\text{max}}$ was sampled from the observed average speed of sc-tracks in the corresponding movie. Likewise, $p_{\text{turn}}(t_p)$ was set based on the empirical "survival" hazard function of persistent intervals in sc-tracks. The size parameter $r_{\text{max}}$ was set to 20 µm, roughly matching observed T cell sizes.

Where it was not possible to derive model parameters from data, we opted for conservative choices that would make it as easy as possible to allow motile trains. If the model then does *not* support motile trains, we can reasonably assume that this is not simply a consequence of the chosen parameters.

For this reason, $r_{\text{min}}$ was set slightly lower at 17, allowing cells to compress slightly but not too much. This choice ensures that particles regain their desired speed relatively quickly after conflict resolution, a conservative choice that makes it as easy as possible for the extended queuing model to support motile T cell trains. Following the same logic, $\tau$ was chosen at 10 steps, which means that particles can regain their desired speed quickly, in less than the time between frames. $\beta$ was set at the same value as used in the original model [1], but given that $\tau$ is small, its value is not too important. Finally, by setting $M = 20s$ equal to the frame rate and $p_{\text{switch}} = 0.5$ at a relatively high value, we ensure that conflicts between particles colliding with opposite $\vec{e}_{\text{target}}$ can be resolved quickly, again making it difficult for traffic jams to form.

**Simulation set-up.** Simulations were constructed with a temporal resolution of one step per second and a duration matched to that of the corresponding movie. Each simulation contained as many 1D lines as the corresponding movie had microchannels, with line length $L$ matched to the width of the field of view in the movie. Particles were initialized with locations $x_0$ and initial direction $\vec{e}_{\text{target}}$ derived from the first two frames of the movie.

**Boundary condition.** An open boundary was implemented by removing the particles once they left the field of view (i.e., $x(t) < 0$ or $x(t) > L$), and letting new particles enter whenever the number of particles in the simulation dropped below the number of cells at the corresponding time in the movie. New particles enter in a random channel from a random side ($x = 0$ or $x = L$); only if there is not already a particle within $d = 2r_{\text{max}}$ of that position.

**Simulation outputs.** Every 20 seconds (= 20 simulation steps), we recorded the centroids of all particles within the field of view, as well as all pairs of particles that were in contact. For this purpose, we considered particles to be in contact if their distance was below 2.5 $r_{\text{max}}$. This threshold is slightly larger than the one used to compute escape velocities since otherwise prolonged "contacts" are by definition not possible in this model. This choice corresponds with the idea that cells are deformable and true physical exclusion does not start immediately upon cell-cell contact, but at slightly smaller distances. Tracks and annotated contacts were then analyzed in the same way as the (preprocessed) T cell data.

#### Fitting CPM motility parameters to a heterogeneous population of sc-tracks.

**Problem definition.** We define cells $c_i = (\lambda_{\text{act}}, \text{max}_{\text{act}})_i$ as heterogeneous individuals sampled from a lognormal distribution: $(\ln \lambda_{\text{act}}, \ln \text{max}_{\text{act}}) \sim \mathcal{N}(\boldsymbol{\mu}, \boldsymbol{\Sigma})$, with means $\boldsymbol{\mu} = (\mu_\lambda, \mu_m)$ and covariance matrix:

$$\boldsymbol{\Sigma} = \begin{bmatrix} \sigma_\lambda^2 & \sigma_{\lambda m} \\ \sigma_{\lambda m} & \sigma_m^2 \end{bmatrix}$$

Thus, a heterogeneous cell population is defined by parameters: $\boldsymbol{\theta} = (\mu_\lambda, \mu_m, \sigma_\lambda, \sigma_m, \sigma_{\lambda m})$. To fit $\boldsymbol{\theta}$ to the observed distributions of speeds and arrest coefficients of sc-tracks, we used Approximate Bayesian Computation (ABC) based on sequential Monte Carlo (SMC) as described by Sisson et al [2]. We outline our implementation below along with the choice in prior and distance function.

**Algorithm.** In SMC-ABC, we iteratively propose candidate parameter sets $\boldsymbol{\theta}_k$, which we only "accept" if some user-defined distance $d(\mathcal{S}_{\theta_k}, \mathcal{X})$ between simulated data at parameters $\boldsymbol{\theta}_k$ (abbreviated $\mathcal{S}_{\theta_k}$) and T-cell sc-tracks ($\mathcal{X}$) is below a given threshold $\epsilon$ (see "distance function" and "distance threshold" below). We start with a lenient threshold $\epsilon$=0.5 and reduce it over time, ensuring that $\mathcal{S}_{\theta_k}$ for accepted parameters gets closer and closer to the target data.

Thus, every SMC-ABC "generation"[1] $g$, we:

1. Sample parameters $\boldsymbol{\theta}_k^g$ from the prior distribution (if $g$ = 1) or from $\boldsymbol{\theta}^{g-1}$ (see "priors and candidate sampling"), keeping only $\boldsymbol{\theta}_k^g$ with positive semi-definite covariance matrix, and:
   (a) Draw $N_c$ = 20 cells $c_i = (\lambda_{\text{act}}, \text{max}_{\text{act}})_i$ from the distribution parametrized by $\boldsymbol{\theta}_k^g$;
   (b) Simulate each $c_i = (\lambda_{\text{act}}, \text{max}_{\text{act}})_i$ individually in a 13 pixel high microchannel with periodic boundaries for a duration $N_{\text{steps}}$ sampled from the empirical sc-tracks, at 0.454 µm/pixel, 0.5 seconds/MCS, and recording centroids every 40 MCS (other model parameters as in Table S4);
   (c) Use the resulting $\mathcal{S}_{\theta_k}$ (of $N_c$ simulated tracks) to compute $d(\mathcal{S}_{\theta_k}, \mathcal{X})$ (see "distance function").
   (d) Accept $\boldsymbol{\theta}_k^g$ if and only if $d(\mathcal{S}_{\theta_k}, \mathcal{X}) < \epsilon$ (i.e. simulated tracks resemble T-cell sc-tracks).
2. Repeat step 1 until we either:
   (a) Have 500 accepted parameter sets $\boldsymbol{\theta}_k^g$ and continue to step 3, or
   (b) Fail to accept any $\boldsymbol{\theta}_k^g$ for 50 consecutive attempts; in which case we stop the algorithm and return the 500 accepted $\boldsymbol{\theta}_k^{g-1}$ from the previous generation;
   (c) Complete 15 generations, in case we also stop the algorithm.
3. Update $\epsilon$ to be the median of the accepted distances $d(\mathcal{S}_{\theta_k}, \mathcal{X})$ in step 2;

We repeat step 1-3 until termination, using the obtained sample from the 5-dimensional posterior to get the maximum a posteriori estimate using kernel density estimation.

**Priors and candidate sampling.** In the first generation, we use uniform priors $\mu_\lambda \sim \mathcal{U}(5, 7)$ , $\mu_m \sim \mathcal{U}(2.5, 4.5)$, $\sigma_\lambda \sim \mathcal{U}(0.2, 0.6)$, $\sigma_m \sim \mathcal{U}(0.2, 0.6)$ to ensure $\text{max}_{act}$ falls in range $[0, 120]$ and $\lambda_{act}$ in range $[0, 1400]$, covering realistic cell motility [3], and $\sigma_{\lambda m} \sim \mathcal{U}(-1, 1)$ to make no prior assumptions on parameter covariance. After the first generation, new candidates $\boldsymbol{\theta}_k^g$ are proposed by mutating accepted $\boldsymbol{\theta}_k^{g-1}$ from the previous generation, adding independent Gaussian noise to each parameter $p \sim \mathcal{N}(0, \sigma_p)$, with $\sigma_p^2$ equal to 1/500th of the prior range of the given parameter (e.g. for $\mu_\lambda$, 1/500 * (7-5) = 0.004).

[1] In ABC literature this is often referred to as "population", but we call it generation here to distinguish it from the heterogeneous population of cells we are simulating.

**Distance function.** For a simulated dataset $\mathcal{S}_{\theta_k}$ of $N_c$ simulated tracks simulated at parameters $\theta_k$, we compute the distance $d(\mathcal{S}_{\theta_k}, \mathcal{X})$ to the T-cell data $\mathcal{X}$ as follows. During the simulation, we check for cell breaking using the "connectedness" defined in [3]; if one of the simulated cells was broken (i.e. the connectedness was less than 95% for more than 1% of the time) the parameter set was automatically rejected (i.e. $d(\mathcal{S}_{\theta_k}, \mathcal{X}) = \infty$). In other cases, we computed the distribution of mean track speeds and arrest coefficients on both datasets, normalized to the maximum observed value in $\mathcal{X}$). Instead of computing the arrest coefficient (AC) as the fraction of "stops" in the track, $AC = N_{\text{stop}}/N_{\text{steps}}$, we instead use AC $= \frac{(N_{\text{stop}}+1)}{(N_{\text{steps}}+2)}$ to prevent zero values, such that we can use the distribution of logit-transformed AC (which increases robustness of the algorithm). We then perform moments matching on the first three moments of speed and logit-transformed AC distributions (crude mean, variance, and skewness), defined for $m$ = 1,2,3 as:

$$E\left[X^m\right] = \sum_i^N x_i^m / n$$

Denoting with $\Delta M_m(\mathcal{S}_{\theta_k}, \mathcal{X})$ the absolute difference in $m$-th moment between simulation and data:

$$d(\mathcal{S}_{\theta_k}, \mathcal{X}) = \frac{1}{2} \sum_{m=1}^{3} w_m^{\text{speed}} \Delta M_m^{\text{speed}}(\mathcal{S}_{\theta_k}, \mathcal{X}) + w_m^{\text{AC}} \Delta M_m^{\text{AC}}(\mathcal{S}_{\theta_k}, \mathcal{X})$$

with weights inversely proportional to the variance of the empirical moments such that moments with lower variance contribute more:

$$w_m^X = \frac{c}{\text{Var}\left(E\left[X^m\right]\right)} = \frac{cn^2}{\sum \left(x_i^m - \mu_m\right)^2}, \quad s.t. \quad \sum_{m=1}^{3} w_m^X = 1$$

Here, constant $c$ is chosen such that the weights sum up to one.

### Supplementary Tables

#### Table S1: Kaplan-Meier estimates of motile T cell train survival.

Kaplan-Meier estimates of median and 95% confidence interval (CI) of motile (>5μm/min) T-cell train survival. Corresponds to Fig. 3d; NA estimates indicate quantiles that could not be estimated due to censoring.

| Train size | Number of trains | Events | Median survival time | 95% CI |
|---|---|---|---|---|
| 2 | 812 | 106 | NA | NA,NA |
| 3 | 489 | 90 | NA | 26 min,NA |
| 4 | 259 | 54 | 12.3 min | 8 min,NA |
| >4 | 436 | 90 | 14.3 min | 9.7 min,NA |

#### Table S2: Kaplan-Meier estimates of motile Neutrophil train survival.

Kaplan-Meier estimates of median and 95% confidence interval (CI) of motile (>5μm/min) Neutrophil train survival. Corresponds to Fig. 6g.

| Train size | Number of trains | Events | Median survival time | 95% CI |
|---|---|---|---|---|
| 2 | 1806 | 829 | 2 min | 1.7 min, 2.3 min |
| 3 | 631 | 322 | 1.3 min | 1.3 min, 1.7 min |
| 4 | 294 | 140 | 1.3 min | 1 min, 1.3 min |
| >4 | 404 | 204 | 1 min | 1 min, 1.3 min |

#### Table S3: Extended queuing model parameter values and descriptions.

| Parameter | Explanation | Value | Justification |
|---|---|---|---|
| $v_i^{\text{max}} = v_{\text{e},i}$ | Desired and escape velocity of particle $i$ | Sampled from observed sc-track speeds (mean 14 µm/min, range ∼5-30 µm/min) | Matching experimental data. |
| $\Delta t$ | Duration of one time step in the model | 1 second | Smaller than $\frac{r_{\text{min}}}{2v_i^{\text{max}}}$ for all $v_i^{\text{max}}$ (see above); as recommended by Baglietto and Parisi[1]. |
| $\vec{e}_{\text{target},i}$ | Target direction of particle $i$ | Initial values based on first frames of movie; then follows from model dynamics | Matching experimental data. |
| $r_{\text{max}}$ | Particle size when moving, affects inst. speed | 20 µm | Similar to observed T-cell sizes. |
| $r_{\text{min}}$ | Particle size in collision | 17 µm | Allow slight, but not extreme compression as qualitatively observed in experimental data . |
| $\tau$ | Relaxation time back to full speed after collision | 10 seconds | Conservative choice; allowing particles to regain their speed quickly makes it easier for the system to stay motile. |
| $\beta$ | Constant affecting relaxation of speed | 0.9 | From original model; but given that $\tau$ is small, has little effect |
| $P_{\text{turn}}(t_p)$ | Probability of flipping the target direction given that the particle has been persistent for time $t_p$ | Extracted from turning hazard function as extracted from T-cell directional "survival" curve | Matching experimental data. |
| $M$, $P_{\text{switch}}$ | If the particle has moved less than 10% of its desired $v_i$ over the last period $M$ due to collisions with other particles, it can flip its direction with probability $p_{\text{switch}}$ | $M$ = 20 seconds, $P_{\text{switch}}$ = 0.5 | Conservative choice; with $M$ relatively low and $P_{\text{switch}}$ relatively high, it is as easy as possible for the particles to stay motile. |
| $L$ | channel width | 625 µm | Matching experimental data. |

#### Table S4: CPM parameter values and descriptions.

| Parameter | Explanation | Value | Justification |
|---|---|---|---|
| $\Delta x, \Delta t$ | Spatial and temporal resolution | 0.454 micron/pixel, 0.5 sec/MCS | Spatial resolution matches images. Temporal resolution balances computational efficiency while still allowing a good fit of motility. |
| $T$ | Parameter controlling stochasticity in the model | $T$ = 5 | Set to lower value compared to published work [3] to have more stable cell behavior upon collision. |
| $J_{\text{cell,bg}}$ | Interface energy of the boundary between the cell and the channel wall or empty medium | $J_{\text{cell,bg}}$ = 0 | Set slightly lower than in published work [3] to simplify interpretation. |
| $J_{\text{cell,cell}}$ | Interface energy of the boundary between two contacting cells | $J_{\text{cell,cell}}$ = $J_{\text{cell,bg}}$ - $\gamma$, where $\gamma$ is the "adhesion" parameter which is zero unless otherwise specified. | If $J_{\text{cell,cell}}$ = $J_{\text{cell,bg}}$ ($\gamma = 0$), cell-cell contact is energetically neutral, which we assume until proven otherwise. |
| $V_{\text{target}}$ | Desired size (in pixels) of the cell | 500 | At a resolution of 0.454 um/pixel, this yields cell sizes that are similar to those observed for T cells. |
| $\lambda_V$ | Parameter controlling the importance of maintaining size | 1 | Allows slight but not strong compressibility. |
| $P_{\text{target}}$ | Desired size of the cell boundary | 310 | Set slightly lower than in published work [3] to compensate for the lower $J_{\text{cell,bg}}$ |
| $\lambda_P$ | Parameter controlling the importance of maintaining boundary size | 2 | From published work [3] |
| $\max_{\text{act}}$ | Stability of polarity; this is how long pixels "remember" their recent protrusive activity (in MCS). | Sampled from lognormal distribution parametrized by $\theta$, see below. | Allow for a population of heterogeneous individuals; fitted to match sc-track speed and arrest coefficient distributions. |
| $\lambda_{\text{act}}$ | Importance of the positive feedback on activity compared to the other forces acting on the cell | Sampled from lognormal distribution parametrized by $\theta$, see below. | Allow for a population of heterogeneous individuals; fitted to match sc-track speed and arrest coefficient distributions. |
| $\eta$ | Force transfer efficiency parameter: which fraction of force $\Delta H_{\text{act}}$ can be propagated in case of copy attempts into another cell? | Varied; $\eta = 1$ corresponds to the usual Act-CPM. | Show the effect by examining the extremes. |
| $\boldsymbol{\theta}$ | $\boldsymbol{\theta} = (\mu_\lambda, \mu_m, \sigma_\lambda, \sigma_m, \sigma_{\lambda m})$ parametrizes a lognormal distribution $(\ln \lambda_{\text{act}}, \ln \max_{\text{act}}) \sim \mathcal{N}(\boldsymbol{\mu}, \Sigma)$ from which each cell's motility parameters are sampled during the simulation. $\boldsymbol{\mu} = (\mu_\lambda, \mu_m)$ and $\Sigma = [[\sigma_\lambda^2, \sigma_{\lambda m}], [\sigma_{\lambda m}, \sigma_m^2]]$ | $\mu_\lambda$ = 5.5763, $\mu_m$ = 4.1242, $\sigma_\lambda^2$ = 0.4653, $\sigma_m^2$ = 0.2882, $\sigma_{\lambda m}$ = -0.0707 | Fitted on T cell sc-track data; see Methods and Supplementary Methods. |

#### Supplementary Video Legends

##### Video S1: T cell traffic in microchannels.

Left: Video corresponding to Fig. 1d (note: Fig. 1d corresponds to the top channels in this video but is mirrored compared to the video.) The T-cell train shown in Fig. 1e is the train at about 25% from the top, starting from 00:05:20 (see also Video S2). The single T cell in Fig. 1e corresponds to the cell that comes into that same channel from the right at around 00:22:00s; the timelapse in Fig. 1e starts at 00:27:40s. Right: Same video, now with extracted sc-tracks (circles) on top.

##### Video S2: A T cell train in a microchannel.

Example T cell train from Fig. 1e, zoomed in from Video S1.

##### Video S3: Single-cell motion.

Videos showing the cells in Fig. 2d and the corresponding raw and smoothed speeds in Fig. 2e. (These cells come from a different movie than Video S1).

##### Video S4: Example collisions.

Head-to-head and head-to-tail collisions from Fig. 3a and Fig. 4a.

##### Video S5: Example simulation of the extended queuing model.

Purple gradient compares the effective speed over the last 10 steps as a percentage of the agent's desired speed. Horizontal lines indicate "contacts" (at slightly larger distances than the agent size as described in Methods). Small purple dots indicate the current direction of the agent.

##### Video S6: Example Cellular Potts model simulation.

Darker gray indicates the "active" protruding region at the front of the cell. Blue dots indicate the center of mass. Corresponding to Fig. 5.

##### Video S7: Neutrophil traffic in microchannels.

An example of neutrophil traffic in microchannels. Corresponding to Fig. 6b.

##### Video S8: Neutrophil queuing and jamming.

Corresponding to Fig. 6e.
